# High fat diet disrupts diurnal interactions between REG3γ and small intestinal gut microbes resulting in metabolic dysfunction

**DOI:** 10.1101/2020.06.17.130393

**Authors:** Katya Frazier, Amal Kambal, Elizabeth A. Zale, Joseph F. Pierre, Nathaniel Hubert, Sawako Miyoshi, Jun Miyoshi, Daina Ringus, Dylan Harris, Karen Yang, Candace Cham, Mark W. Musch, Eugene B. Chang, Vanessa Leone

**Affiliations:** Department of Medicine, The University of Chicago, Chicago, IL 60637; Infectious Diseases Division, Weill Cornell Medicine, New York, NY, 10065; Department of Pediatrics, University of Tennessee Health Science Center, Memphis, TN 38163; Department of General Medicine, Kyorin University School of Medicine, Tokyo, Japan 1818611; Department of Gastroenterology and Hepatology, Kyorin University School of Medicine, Tokyo, Japan 1818611; Northwestern University Feinberg School of Medicine, Northwestern University, Chicago, IL 60611

**Keywords:** Circadian rhythms, Innate immunity, *Reg3γ*, Gut microbiota, Small intestine, High-fat diet, Diurnal oscillation, Organoid, Germ-free, Host-microbe interactions

## Abstract

Gut microbial diurnal oscillations are important diet-dependent drivers of host circadian rhythms and metabolism that ensure optimal energy balance. Yet, the interplay between diet, microbes, and host factors that sustain intestinal oscillations is complex and poorly understood. Here, we report the host C-type lectin antimicrobial peptide *Reg3γ* works with key ileal microbes to orchestrate these interactions in a bi-directional manner, independent from the intestinal core circadian clock. High fat diet diminishes physiologically relevant microbial oscillators essential for host metabolic homeostasis, resulting in arrhythmic host *Reg3γ* expression and increased abundance and oscillation of *Reg3γ*-independent gut microbes. This illustrates a transkingdom co-evolved biological rhythm involving reciprocating, sensor-effector signals between key host and microbial components that ultimately drive metabolism, but are also heavily influenced by diet. Restoring the gut microbiota’s capacity to sense and transduce dietary signals mediated by specific host factors such as *Reg3γ* could be harnessed to improve metabolic dysfunction.

## Introduction

Disruption of circadian rhythms (CRs) in modern society has contributed to the rise in incidence of metabolic diseases (Turek et al., 2005). CRs are 24-hr oscillations in behavioral and biological processes driven in part by a core circadian clock (CC) transcriptional-translational feedback loop found in nearly all cells within the body (Dibner et al., 2010). These rhythms are crucial to regulation of major metabolic and immune pathways, where nearly half the murine transcriptome is under CC control (Zhang et al., 2014). When CRs are altered or non-functional, adverse metabolic consequences emerge, including increased adiposity and impaired insulin sensitivity (Turek et al., 2005; Jacobi et al., 2015; Karlsson et al., 2003).

Recent work shows the trillions of gut microbes not only influence host digestion, absorption, and energy balance, but are intricately intertwined with the host CC, where disruption either via environmental manipulations, (Thaiss et al., 2014), high fat diet (Leone et al., 2015; Zarrinpar et al., 2014) or via genetic mutation (Liang et al., 2015; Thaiss et al., 2016), results in loss of gut microbiome oscillations. Diurnal microbiome oscillations in cecum, colon, and stool are driven, in part, by time of feeding and nutrient delivery to the gut over 24hrs. However, we previously revealed mice fed via continuous parenteral nutrition (PN) drives unique gut microbial community membership, yet PN microbiota still exhibited diurnal oscillations (Leone et al., 2015). This suggests additional gut signals, whether host-derived, microbially-derived, or both, can play a role in gut microbial rhythmicity.

While host factors drive microbial rhythms, microbes provide feedback that influence host diurnal patterns both locally and peripherally. In mice, gut microbes drive rhythmicity and amplitude of CC and innate immune factors in intestinal epithelial cells (IEC) that influence host lipid metabolism (Wang et al., 2017). Gut microbe antibiotic depletion or their complete absence, i.e. germ-free (GF), significantly impairs circadian dynamics of IEC Toll-like receptor (TLR) expression and downstream gene that drive intestinal antimicrobial peptide (AMP) synthesis, a group of host-derived molecules that target specific microorganisms to aid in maintaining gut homeostasis (Mukherjee and Hooper, 2015).

One particular AMP associated with host-microbe diurnal patterns is *Regenerating islet-derived protein 3 gamma* (*Reg3γ*). REG3γ is a Myeloid differentiation primary response 88 (MyD88)-dependent C-type lectin made by IECs throughout the GI tract, and highly expressed in the distal small intestine (SI) (Narushima et al., 1997; Nata et al., 2004; Vaishnava et al., 2008) that targets Gram-positive bacteria (Cash et al., 2006a; Mukherjee et al., 2014). REG3γ aids in separating IECs from mucosa-associated microbes, while diet-induced obesity (DIO) ablates *Reg3γ* expression (Everard et al., 2014; Loonen et al., 2014). Antibiotic depletion of gut microbes significantly impairs diurnal *Reg3γ* expression in colon and ileum (Mukherji et al., 2013). *Reg3γ*-deficient mice exhibit loss of host-microbial separation (Vaishnava et al., 2011; Wang et al., 2016) and disrupted diurnal rhythms of mucosa-associated microbial abundance within the colon (Thaiss et al., 2016). How diurnal *Reg3γ* expression aids in maintaining normal gut microbial oscillations and the implications for host metabolic health remain unexplored.

We investigated whether signals derived from specific gut bacteria are required to drive host diurnal expression patterns of *Reg3γ,* and if these endogenous cues can “reset” bacterial oscillations. We aimed to identify whether microbial signals were lost in high fat diet (HF)-induced gut dysbiosis, further exacerbating host metabolic disruption. Using both *in vivo* and *in vitro* approaches, we show distal SI *Reg3γ* diurnal expression is not under CC control, but is regulated locally by specific, diet-dependent oscillating bacteria. We reveal small molecules derived from bacteria promoted by regular chow diet (RC), but not HF, induce *Reg3γ* expression *in vitro*, and these bacteria exhibit unique susceptibility to REG3γ’s antimicrobial action. Finally, we reveal *Reg3γ* deficiency coupled with HF permits a gain in oscillation in relative abundances of specific gut microbes normally susceptible to REG3γ. Together, our data demonstrate a reciprocating, synchronized relationship between rhythmic, diet-selected SI gut microbes and REG3γ, where desynchronization of these local interactions can result in metabolic disruption and perhaps DIO.

## Results

### Diurnal expression of SI *Reg3γ* is independent from the core CC gene network and requires diet-induced gut microbiota

We first examined impact of diet and microbe status on core CC genes within distal ileum mucosal scrapings (DIMS) obtained from GF and SPF mice fed RC or HF. We observed some significant, yet modest core CC gene expression changes within specific Zeitgeber time points (ZT), i.e., *Bmal1*, *Clock*, and *Cry1* over a 12:12 LD cycle (**Figure 1A, Figure S1A**). Despite changes within ZT, neither RC, HF, or gut microbe status impacted diurnal rhythmicity and amplitude of core CC genes in DIMS (**Table S1**).

**Figure 1.**
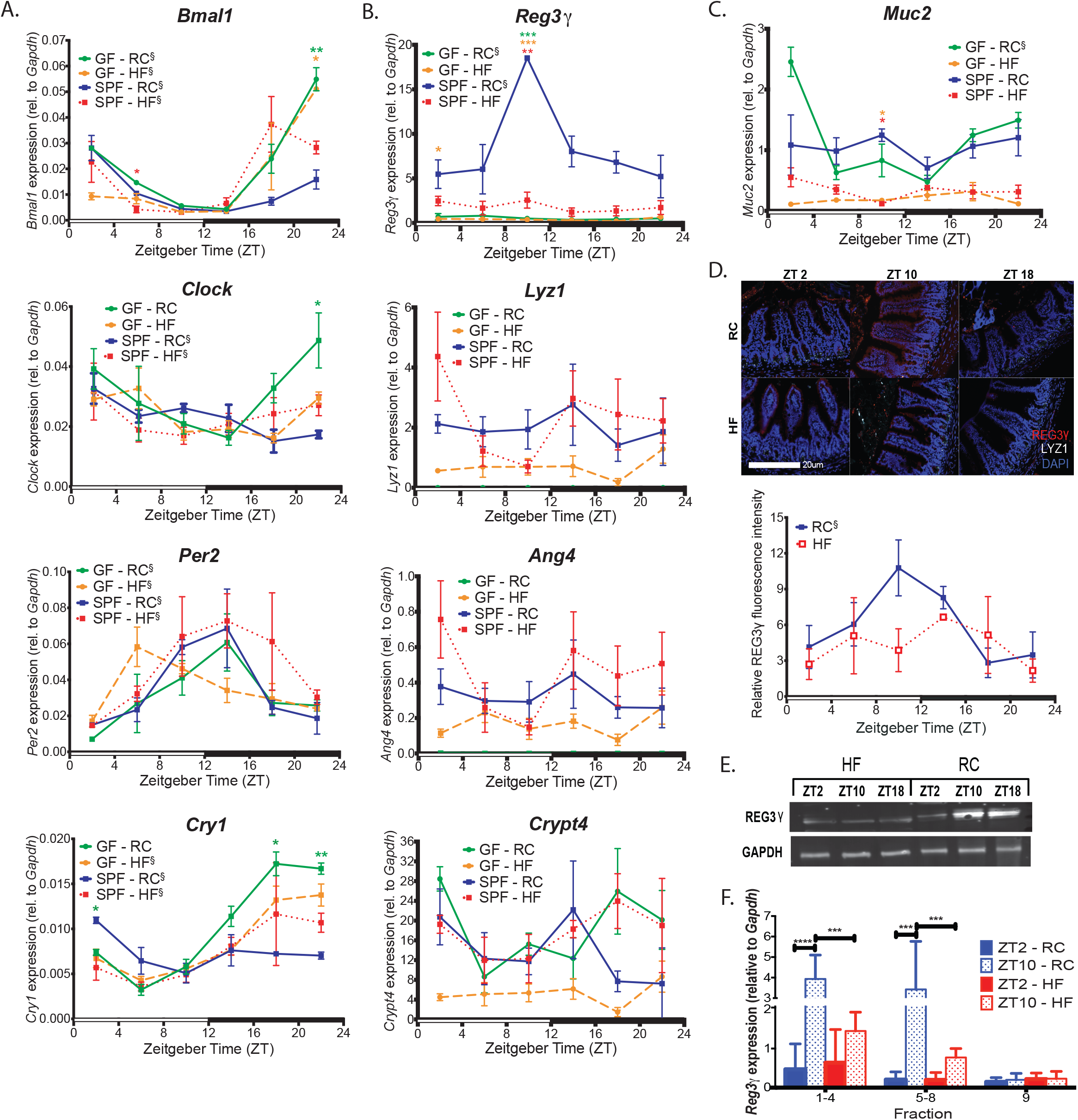
Diurnal patterns of SI *Reg3γ* expression is independent of the core CC network and requires presence of diet-induced gut microbes. **(A - C)** Circadian (A), anti-microbial peptide (B), and *Mucin*2 (C) gene expression in DIMS collected every 4hrs from RC or HF-fed GF and SPF mice maintained in 12:12 LD (indicated by open and closed bars on the X-axis). *p<0.05; **p<0.01; ***p<0.001 via Brown-Forsythe and Welch ANOVA followed by Dunnett’s test as compared to SPF-RC control—star color indicates treatment exhibiting significance within ZT. ξ indicates significant (p<0.05) co-sinor expression patterns detected via CircWave. **(D)** Representative immunostaining images for REG3γ in distal ileum sections from RC and HF-fed SPF mice at ZT 2, 10, and 18, with corresponding fluorescence intensity quantification. Welch’s t test was performed within ZT. **(E)** REG3γ Western blot in DIMS from RC and HF-fed SPF mice at ZT2, 10, and 18. **(F)** *Reg3γ* expression in distal ileum epithelial fractions from RC or HF-fed SPF mice at ZT2 and 10. Fractions 1-4 and 5-8 = absorptive enterocytes, fraction 9 = crypts harboring stem cells and Paneth cells. ***p<0.001; ****p<0.0001 via two-way ANOVA followed by Tukey’s test.

We next examined diurnal transcript levels of several highly expressed AMPs in DIMS of RC or HF-fed SPF and GF mice. RC-fed SPF mice exhibited significantly increased transcript levels within specific ZTs relative to all other groups, particularly *Reg3γ* (**Figure 1B**). Additional AMPs, including β-1,4-glycosidase *Lysozyme1 (Lyz1*), ribonuclease *Angiogenin4 (Ang4)*, and α-defensin *Cryptidin4 (Crypt4*) did not exhibit significant differences between groups. Only *Reg3γ* exhibited diurnal rhythmicity in SPF and GF RC-fed mice, despite significantly reduced overall expression and amplitude in GF animals (**Table S1**). Peak *Reg3γ* expression in SPF RC-fed mice was observed at ZT10, 2hrs prior to lights off and presumably prior to onset of feeding. *Mucin-2* (*Muc2*) expression, which has been previously associated with *Reg3γ* (Huang et al., 2017), was also significantly elevated in RC-fed SPF mice at ZT10 (**Figure 1C**). Only RC-fed GF mice exhibited diurnal *Muc2* expression, although overall levels were not changed relative to SPF RC-fed mice.

We confirmed *Reg3γ* diurnal patterns were also evident at the translational level. Similar REG3γ diurnal patterns were observed in SPF RC-fed, but not in SPF HF-fed counterparts via immunostaining (**Figure 1D,E**). Further analysis of DIMS from SPF RC and HF-fed mice via Western blot (**Figure 1F**) revealed both diet and ZT-dependent REG3γ oscillations. Since *Reg3γ* is expressed by both absorptive enterocytes and Paneth cells (Cash et al., 2006a; Narushima et al., 1997; Nata et al., 2004), we next determined if diurnal and diet-dependent *Reg3γ* expression was lost in all compartments following HF. We performed mucosal villus to crypt cell fractionation using timed EDTA exposure of the gut epithelium, followed by qPCR for *Reg3γ* at ZT2 and 10, the nadir and peak of expression. Pooled fractions 1-8 contained absorptive enterocytes and fraction 9 contained crypts (including Paneth cells). We first confirmed cell fraction composition via marker genes *sucrase-isomaltase* (villus epithelial marker) and *Lyz1* (Paneth cell marker) (**Figure S1B**). Only fractions 1-8 from RC-fed mice exhibited ZT-dependent *Reg3γ* expression, while crypt expression was not different between ZT2 and 10 in RC or HF-fed mice. Together, these data show the core CC gene network is stable regardless of microbe status or diet; however, diurnal rhythms of host *Reg3γ* expression within the villous epithelium are driven by both presence of gut microbes and those selected by diet.

### HF alters ileal microbiota community membership and dampens microbial diurnal oscillations relative to RC

We examined microbial community membership within distal ileal luminal contents (DILC) collected every 4hrs from RC or HF-fed SPF mice via 16S rRNA gene amplicon sequencing and QIIME. HF significantly increased relative abundance of Operational Taxonomic Units (OTUs) belonging to phyla Firmicutes and decreased Bacteroidetes (**Figure 2A**). No differences were detected in less abundant phyla (**Figure S2A**). Beta-diversity analysis using Bray-Curtis (**Figure 2B**, Principal Coordinate Analysis (PCoA)) and Canberra distances (**Figure S2B**) revealed significant differences between RC and HF DILC microbial communities (**Table S3**).

**Figure 2.**
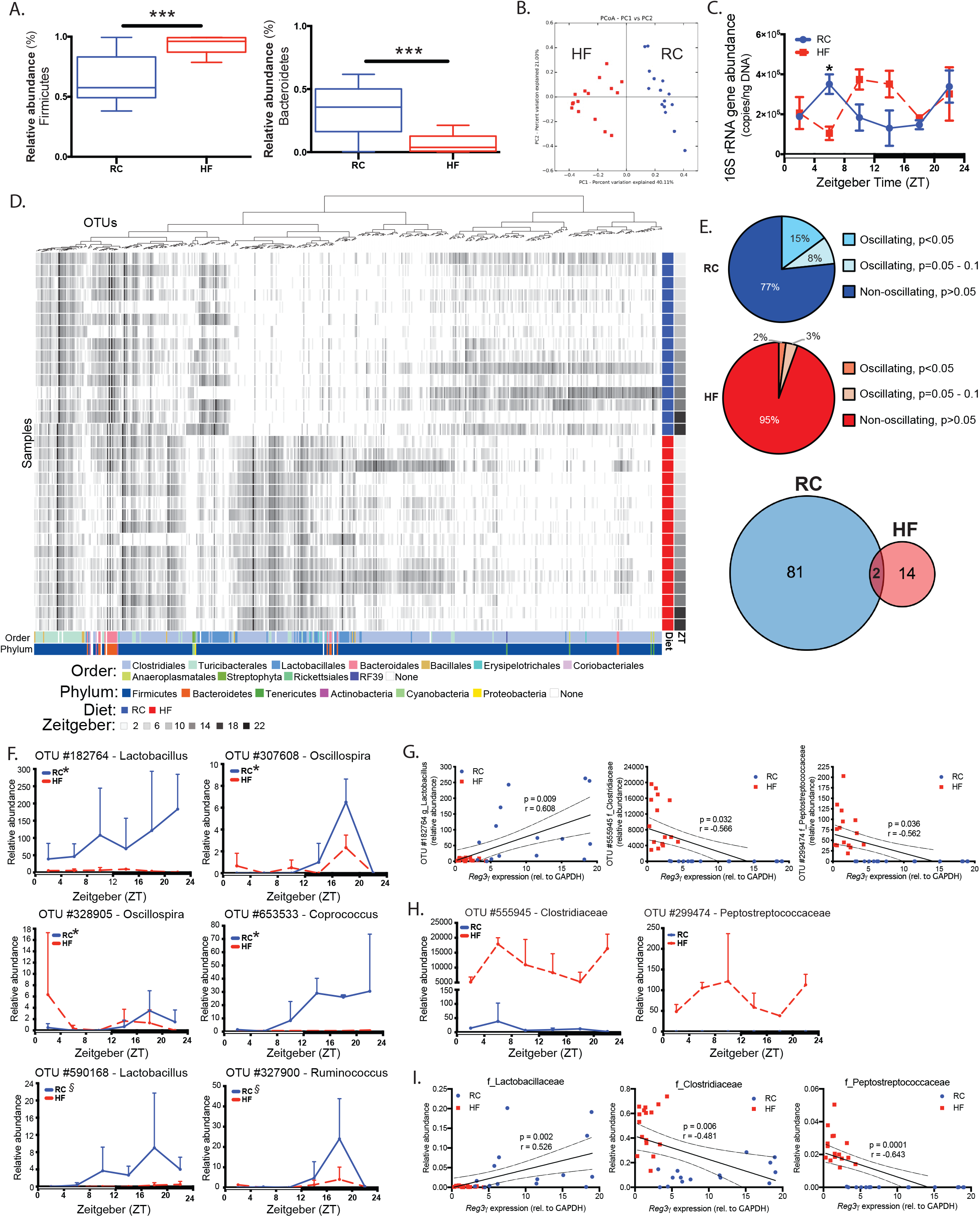
Diet shapes distal SI gut microbe community membership and oscillations that correlate with *Reg3γ* expression. **(A)** Relative abundances of dominant phyla in DILC from RC or HF-fed SPF mice via 16S rRNA gene amplicon sequencing averaged across all timepoints within diet treatment. Box-whisker plots represent mean±min/max. ***p<0.001 via Welch’s t test. **(B)** Bray-Curtis PCoA of 16S rRNA amplicon sequences from RC (blue) and HF (red)-fed mice. **(C)** Absolute 16S rRNA gene copy number determined via qPCR in RC or HF-fed mice. Data represent mean±SEM. *p<0.05 via Welch’s t test. **(D)** anvi’o heatmap of 16S rRNA relative abundances in DILC from RC or HF-fed mice (*n*=2-3 mice/ZT). Columns = OTUs, rows = samples. Colored bars at the bottom represent taxonomy. Blue and red bars to the right represent diet, gray bars represent ZT. **(E)** Percentage (pie charts) of oscillating and non-oscillating OTUs determined via eJTK and numbers (venn diagram) of unique and shared oscillators in DILC from mice fed RC or HF. **(F)** Relative abundances of significantly oscillating OTUs annotated to genus only in RC-fed mice. Data represent mean±SEM. *indicates significant oscillation (p<0.05); ξindicates significant oscillation (p=0.05 − 0.1) detected via eJTK. **(G)** *Reg3γ* expression vs. relative abundances of OTUs exhibiting significant Pearson correlations (Bonferroni p<0.05) in RC and HF-fed mice. Linear regression lines with 95% confidence bands shown. **(H)** Relative abundance of OTUs exhibiting significant negative correlation with *Reg3γ* expression in RC and HF-fed mice. **(I)** *Reg3γ* expression vs. relative abundances of OTUs at the family level that exhibit significant Pearson correlations in RC and HF-fed mice. Linear regression lines with 95% confidence bands shown.

Absolute 16S rRNA gene copy number determined via qPCR averaged across ZT in DILC was not different between RC and HF (**Figure S2C**). Significant differences were only observed between RC and HF-fed mice at ZT6, while diurnal rhythmicity was not detected in RC or HF-fed mice. However, while 16S rRNA gene copy number exhibited a similar amplitude in RC and HF-fed mice, HF elicited a phase-shift relative to RC (**Figure 2C**). Distinct DILC OTU differences were observed between RC and HF-fed mice (**Figure 2D**), where HF significantly increased Clostridiales and decreased Bacteroidales relative to RC (**Table S2**).

To determine whether RC or HF impacted DILC diurnal oscillations, we applied empirical JTK-Cycle (eJTK) to relative abundance data as previously described (Leone et al., 2015). Similar to cecum and feces, ~15% of DILC taxa oscillated under RC and was dramatically reduced by HF (**Figure 2E**). 81 unique oscillating OTUs were observed in DILC of RC-fed mice, while 14 unique OTUs were present in HF-fed mice; only two oscillating OTUs overlapped in RC and HF (**Table S4**). Oscillating RC OTUs annotated to genus were absent, decreased in overall abundance, or lost oscillation under HF (**Figure 2F**). Despite decreased percentage of oscillating OTUs in HF, several gained oscillations (**Figure S2D**). Two OTUs exhibiting oscillations under both RC and HF were annotated to genera *Ruminococcus* and *Oscillospira*, exhibiting diet-dependent effects on their overall abundances (**Figure S2E, Table S4**). Together, these data reveal HF dramatically alters distal ileum microbial membership that corresponds to an overall decrease as well as a unique set of oscillating taxa relative to RC.

### Specific taxa promoted by RC or HF correlate with distal SI diurnal *Reg3γ* expression

Using Pearson correlation analysis, we determined only 3 OTUs in DILC of RC and HF-fed mice significantly correlated with *Reg3γ* expression. A *Lactobacillus* OTU positively correlated with *Reg3γ* expression, while Clostridiaceae and Peptostreptococcaceae OTUs negatively correlated (**Figure 2G**). *Lactobacillus* OTU exhibited oscillation only in RC, with dramatically reduced overall levels in HF-fed counterparts at all ZTs (**Figure 2F**, top left panel). Surprisingly, while OTUs negatively correlated with *Reg3γ* did not exhibit significant oscillations in either RC- or HF-fed mice, their relative abundances were significantly increased across all ZT in HF relative to RC (**Figure 2H**). These associations were also apparent at a higher taxonomic classification level, where relative abundances of families Clostridiaceae, Lactobacillaceae, and Peptostreptococcaceae exhibited nearly identical positive and negative correlations with *Reg3γ* expression (**Figure 2I**). Lactobacillaceae was increased during the mid to late dark cycle only in RC, while Clostridiaceae and Peptostreptococcaceae were significantly increased at several ZTs in HF-fed counterparts (**Figure S2F**). These data reveal diet drives unique SI microbiota membership, serving as the primary driver of microbial oscillations. Lactobacillaceae are enriched by RC and exhibit oscillations, while certain OTUs positively correlate with host diurnal *Reg3γ* expression. Conversely, HF diminishes microbial oscillations and promotes Clostridiaceae and Peptostreptococcaceae overall expansion, which negatively correlate with *Reg3γ* expression.

### Complex gut microbiota or individual bacterial strains induced by diet directly influence Reg3γ expression *in vitro*

We examined the direct impact of diet-induced microbial communities on *Reg3γ* mRNA *in vitro* using 3-dimensional intestinal enteroids derived from distal SI of GF or SPF WT mice. We first used enteroids derived from GF WT mice due to their naïve state and nearly undetectable *Reg3γ in vivo.* DILC lysate derived from RC-fed SPF mice harvested at ZT10 (peak of host *Reg3γ* expression, **Figure 1B**) significantly induced *Reg3γ* expression in GF enteroids compared to HF DILC lysate (p<0.05, **Figure 3A**) after 24hr exposure, indicating diet-induced gut microbiota directly and differentially impact *Reg3γ*.

**Figure 3.**
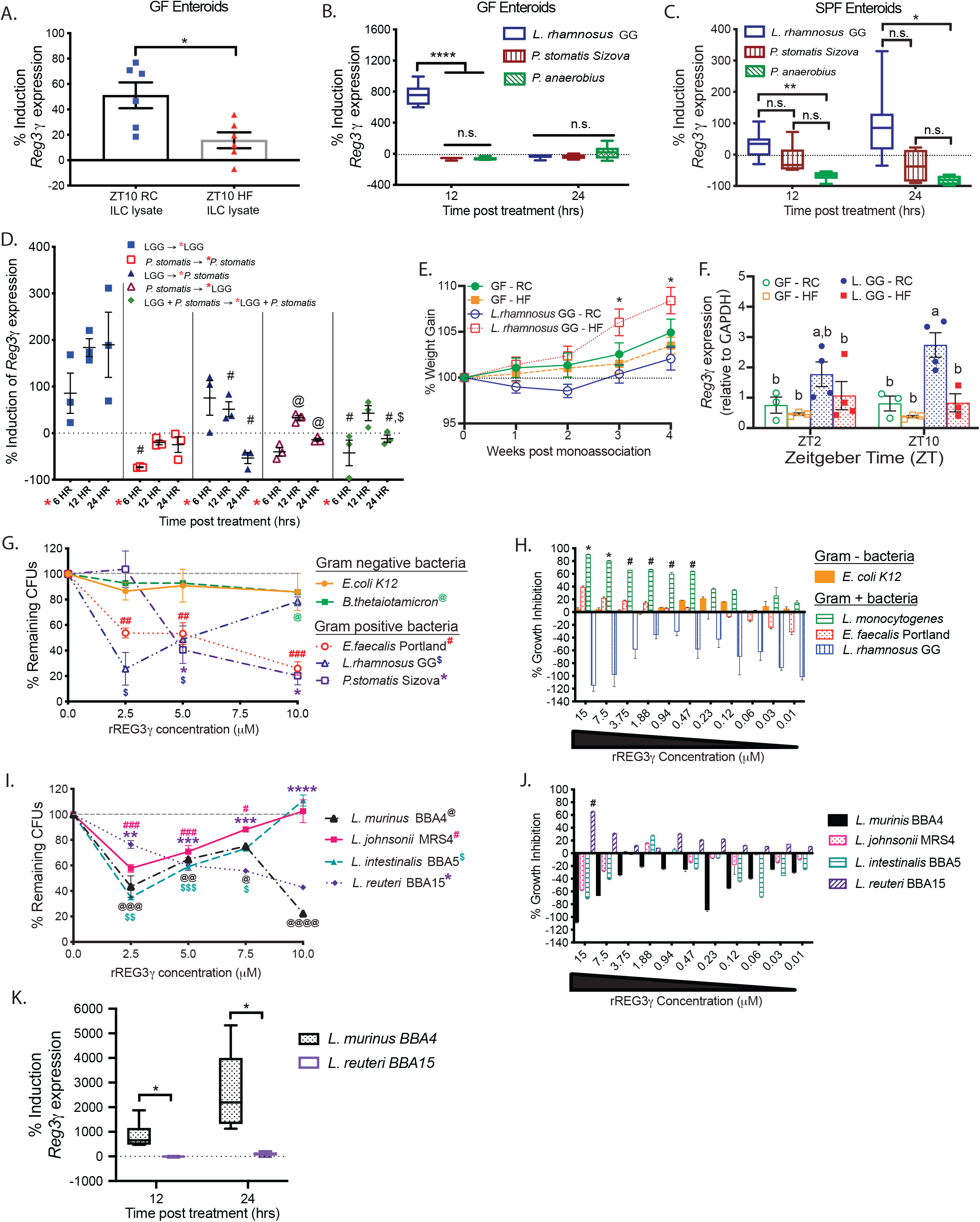
Diet-induced Gram-positive bacteria drive host *Reg3γ* expression and exhibit unique susceptibility to REG3γ’s antimicrobial properties. **(A)** *Reg3γ* expression in GF WT enteroids following 24hr exposure to DILC lysate obtained from RC or HF-fed SPF WT mice at ZT10 relative to PBS vehicle control. *n*=6 technical replicates/treatment. Data representative of 3 independent experiments. Data represent mean±SEM. *p<0.05 via Welch’s t test. **(B,C)** *Reg3γ* expression in GF (B) or SPF (C) WT enteroids following 12 or 24hr exposure to conditioned media (CM) from cultured bacteria strains relative to blank media. *n*=3-6 technical replicates/treatment. Box-whisker plots represent mean±min/max, representative of 3 independent experiments. ****p<0.0001, **p<0.01, *p<0.05 via Brown-Forsythe and Welch ANOVA followed by Dunnett’s test within timepoint; n.s.=not significant. **(D)** *Reg3γ* expression in SPF WT enteroids following 6, 12 or 24hr exposure to CM from cultured bacteria strains relative to blank media. After 6hrs, enteroids were either collected or exposed to second CM treatment. *n*=3 technical replicates/treatment. Data represent mean±SEM, representative of 2 independent experiments. Symbols indicate significant differences relative to: #-12hr LGG; @-6hr *P. stomatis*; $-12hr *P. stomatis* → LGG. p<0.05 via 2-way ANOVA followed by Tukey’s test. **(E)** Percent weight change relative to baseline of RC or HF-fed GF or LGG-monoassociated WT mice. *n*=6-8 mice/treatment. Data represent mean±SEM. *p<0.05 via Brown-Forsythe and Welch ANOVA followed by Dunnett’s test. **(F)** *Reg3γ* expression in DIMS of RC or HF-fed GF or LGG-monoassociated WT mice collected at ZT2 and 10. Data represent mean±SEM, representative of 2 independent experiments. Bars with different letters are significantly different (p>0.05) via two-way ANOVA followed by Tukey’s test. **(G,I)** Percent remaining CFUs of representative (G) or indigenous *Lactobacillus* strains (I) following 2hr exposure to rREG3γ relative to vehicle control. *n*=2 replicates/bacteria strain. Data represent mean±stdev, representative of 3 independent experiments. Symbols indicate significant differences relative to vehicle control; 1 symbol=p<0.05; 2 symbols=p<0.01; 3 symbols=p<0.001; four symbols=p<0.0001, via one-way ANOVA followed by uncorrected Fisher’s LSD. **(H,J)** Minimum inhibitory concentration (MIC) assay of representative (H) or indigenous *Lactobacillus* strains (J) exposed to varying concentrations of rREG3γ. *n*=2 replicates/strain. Data represent mean±stdev, representative of 3 independent experiments. *represents 80% growth inhibition (MIC_80_); #represents 50% growth inhibition (MIC_50_). **(K)***Reg3γ* expression in enteroids derived from SPF WT mice following 12 or 24hr exposure to CM from *L. murinis* BBA4 and *L. reuteri* BBA15. *n*=6 technical replicates/treatment. Data representative of 1 independent experiment. Box-whisker plots represent mean±min/max. *p<0.05 via Welch’s t test.

Next, we determined if bacteria type strains belonging to families significantly correlated with *Reg3γ in vivo* (**Figure 2G,H**) could directly influence expression *in vitro*. Conditioned media (CM) or bacterial lysates were prepared from *Lactobacillus rhamnosus* GG (LGG, Lactobacillaceae), *Peptostreptococcus anaerobius* (Clostridiaceae), and *Peptoanaerobacter stomatis* Sizova (*P. stomatis*, Peptostreptococcaceae,). Enteroids derived from GF WT mice exhibited significant induction of *Reg3γ* expression following 12hr exposure to either LGG-derived CM or lysate (**Figure 3B, S3A**). Neither *P. anaerobius* nor *P. stomatis* CM or lysate significantly induced *Reg3γ* expression. LGG CM similarly induced *Reg3γ* in enteroids derived from SPF WT mice, albeit with a 12hr delay after 24hr exposure (**Figure 3C**). These results indicate that CM is equally capable of *Reg3γ* upregulation on GF or SPF-derived enteroids; therefore, we used SPF-derived enteroids going forward.

We next tested if CM from a non-inducer could inhibit inductive influences of LGG on *Reg3γ* expression. We observed LGG alone induced *Reg3γ* expression, while *P. stomatis* alone resulted in no induction (**Figure 3D**). However, even in the presence of LGG CM, co-exposure to *P. stomatis* CM at any time point suppressed induction of *Reg3γ* expression. These data suggest small molecules or components derived from bacteria exhibiting correlations with *Reg3γ in vivo* directly influence expression in a bacteria species-specific manner.

To test whether CM from LGG or *P. stomatis* requires MyD88 for *Reg3γ* induction as previously shown (Natividad et al., 2013; Vaishnava et al., 2008), enteroids were derived from SPF MyD88^+/−^ or MyD88^−/−^ littermate mice. LGG CM significantly induced *Reg3γ* in *MyD88*^+/−^, but not *MyD88*^−/−^ enteroids, implying LGG requires MyD88, while no induction was observed with *P. stomatis* CM regardless of MyD88 status (**Figure S3B**). CM size fractionation revealed small molecules less than 3kDa derived from LGG were sufficient to induce *Reg3γ* expression, while the suppressive effect of *P. stomatis* was lost in all fractions containing small molecules less than 30kDa (**Figure S3C**). This suggests LGG-derived small molecules and components induce, while *P. stomatis*-derived large molecules and components suppress *Reg3γ* expression in a MyD88-dependent manner.

### *Lactobacillus* ZT-dependent distal SI *Reg3γ* expression is mediated by diet composition

To determine whether our *in vitro* findings translated *in vivo*, we monoassociated GF mice fed either RC or HF with LGG. HF+LGG resulted in a modest, yet significant, body weight increase compared to RC+LGG at 3 and 4 wks post-association (**Figure 3E**). Liver weights were not different, however both GF HF and RC + LGG mice exhibited significantly increased gonadal fat relative to GF RC (**Figure S3D**, top/middle panels). GF HF and HF+LGG exhibited significantly increased mesenteric fat weight relative to GF RC and RC+LGG (**Figure S3D**, bottom panel). *Reg3γ* expression was significantly increased in DIMS only from RC+LGG, but not HF+LGG mice harvested at ZT2 and 10 (nadir and peak expression in RC SPF mice), where ZT10 tended to be higher (**Figure 3F**). *Lyz1* expression was increased only in RC+LGG mice (**Figure S3E**). Together, these data suggest microbes and diet combined impact distal ileal *Reg3γ* with corresponding changes in host metabolic outcomes.

### Diet-induced Gram-positive bacteria are uniquely susceptible to REG3γ

Our observations revealed specific Gram-positive bacteria directly induce or suppress ileal *Reg3γ* expression. To elucidate bidirectionality of this dynamic, we determined if REG3γ exhibited differential bactericidal action against these strains using murine recombinant REG3γ (rREG3γ). We recapitulated previous findings showing Gram-negative bacteria were not susceptible to rREG3γ, whereas only ~30% of Gram-positive *Enterococcus faecalis* Portland CFUs remained at 10μM (Cash et al., 2006a) (**Figure 3G**). Similar to *E. faecalis*, *P. stomatis* CFUs dramatically declined at 5 and 10μM rREG3γ; however, LGG appeared susceptible at low and resistant at high concentrations, suggesting unique susceptibility of representative diet-induced gut bacteria correlating with host *Reg3γ*.

We next determined if LGG’s lack of susceptibility to high rREG3γ concentrations was due to the bactericidal assay’s nutrient-limited conditions and identified rREG3γ’s minimal inhibitory concentration (MIC) in actively dividing bacteria. Following an overnight exposure of Gram-positive and −negative bacteria to rREG3γ concentrations ranging from 15uM to 0.01uM optical density was measured (**Figure 3H**). Relative to vehicle control, *E. coli K12* was unaffected by rREG3γ (0% inhibition), while *Listeria monocytogenes* growth was inhibited by ~80% or greater at 7.5 and 15μM rREG3γ. *E. faecalis* growth was inhibited by ~40% at 15μM rREG3γ and decreased in a dose-dependent manner. LGG growth was not inhibited and appeared to be enhanced at nearly all rREG3γ concentrations. Together, these data suggest *Lactobacillus* may be resistant to REG3γ’s antimicrobial properties relative to other Gram-positive bacteria.

### Murine indigenous distal SI *Lactobacillus* species are uniquely resistant to REG3γ

To this point, our *in vitro* studies utilized bacteria type strains which may not fully recapitulate those found in the murine gut, i.e., LGG was originally isolated from human GI tract. We tested whether our observation regarding LGG was generalizable across *Lactobacillus* in murine-derived strains. We isolated four indigenous *Lactobacillus* strains from DILC of RC-fed WT mice. We revealed *L. johnsonii* MRS4 and *L. intestinalis* BBA5 behaved similar to LGG following rREG3γ exposure (**Figure 3I**). While *L. murinus* BBA4 was nearly identical except at 10μM rREG3γ, *L. reuteri* BBA15 displayed similar susceptibility as *E. faecalis* and *P. stomatis.* The MIC assay also showed *L. johnsonii*, *L. intestinalis*, and *L. murinus* were comparable to LGG, while *L. reuteri* growth was inhibited by ~70% at high rREG3γ concentrations (**Figure 3J**). These data show some, but not all, RC-promoted *Lactobacillus* can resist or evade REG3γ, which could impact their ability to thrive under fluctuating diurnal intestinal conditions.

### REG3γ deficiency promotes diet-independent glucose intolerance and does not impact core CC gene expression in distal SI

We next examined how host REG3γ deficiency impacts host and microbial dynamics in RC and HF-fed SPF global *Reg3γ* knockout (*Reg3γ*^−/−^) and heterozygote littermate control (*Reg3γ*^+/−^) mice. *Reg3γ*^−/−^ mice were equally susceptible to HF DIO relative to *Reg3γ*^+/−^ littermates, where body weight increased ~12% relative to RC-fed counterparts over 4 weeks (**Figure 4A**). Liver weights were not different while HF significantly increased gonadal and mesenteric fat relative to RC regardless of genetic background (**Figure S4A**). RC-fed *Reg3γ*^−/−^ mice exhibited significantly worse glucose tolerance, with slower glucose clearance rates and increased area under the curve (AUC) compared to RC-fed *Reg3γ*^+/−^ controls while HF decreased blood glucose clearance rate regardless of genetic background relative to RC-fed *Reg3γ*^+/−^ mice (**Figure 4B**). These data suggest REG3γ does not alter HF DIO, but rather plays a functional role in how diet impacts glucose tolerance, where REG3γ deficiency impairs glucose clearance rates regardless of diet.

**Figure 4.**
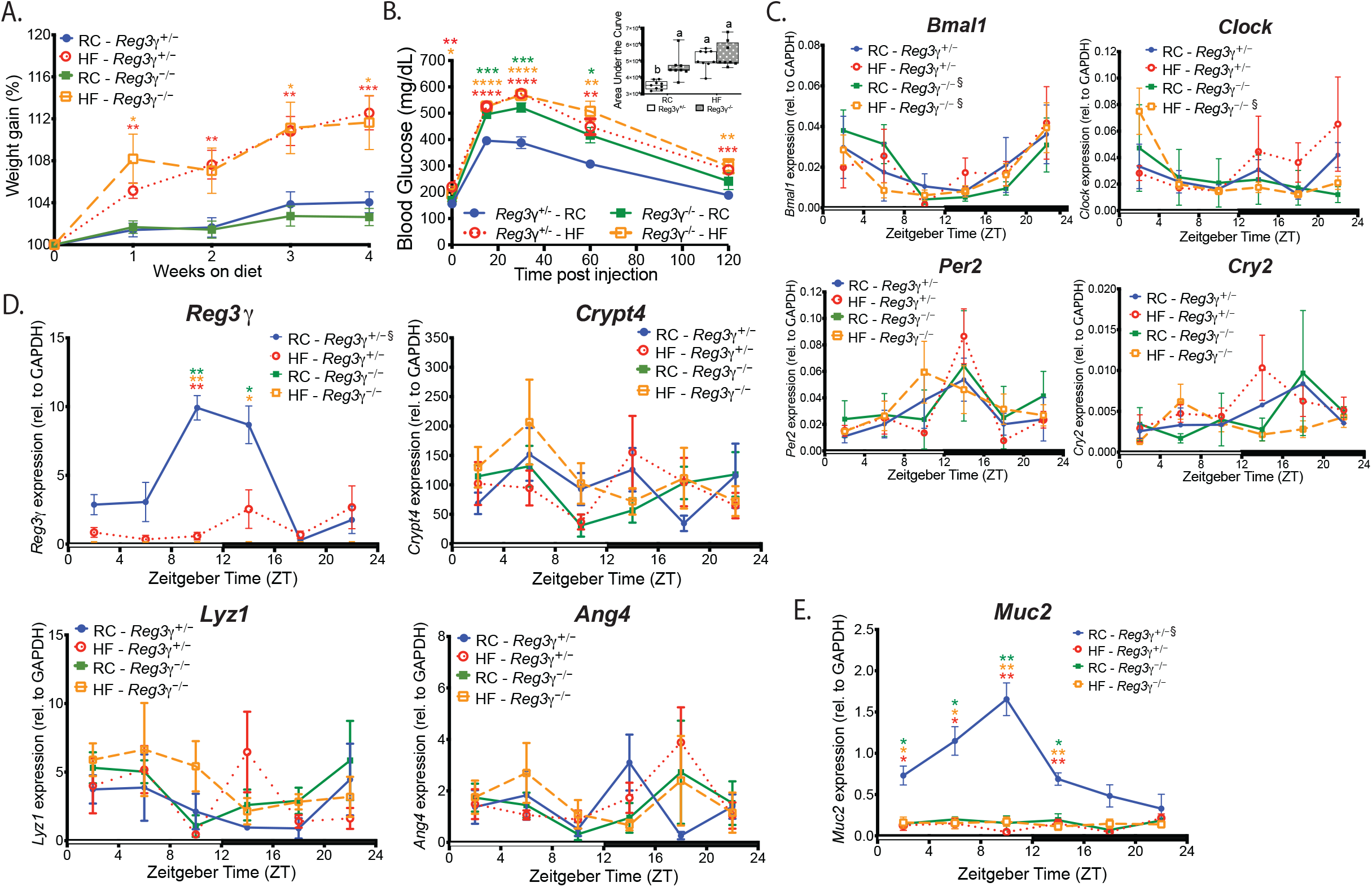
*Reg3γ* deficiency impacts glucose homeostasis in a diet-dependent manner independent from the core CC network. **(A)** Percent weight change from baseline in RC or HF-fed *Reg3γ*^+/−^ or *Reg3γ*^−/−^ mice fed. *n*=23-25 mice/treatment. Data represent mean±SEM. *p<0.05; **p<0.01; ***p<0.001 via Brown-Forsythe and Welch ANOVA followed by Dunnett’s test as compared to RC-fed *Reg3γ*^+/−^ control—star color indicates treatment exhibiting significance within time point. **(B)** Glucose tolerance test in *Reg3γ*^+/−^ or *Reg3γ*^−/−^ mice fed RC or HF. *n=*8-9 mice/treatment. Data represent mean±SEM. *p<0.05; **p<0.01; ***p<0.001; ****p<0.0001 via Brown-Forsythe and Welch ANOVA followed by Dunnett’s test relative to *Reg3γ*^+/−^ RC control—star color indicates treatment exhibiting significance within time point. Inset graph represents Area Under the Curve. (AUC). Box-whisker plots represent mean±min/max. Bars with the same letter are not significantly different (p>0.05) via two-way ANOVA followed by Tukey’s test. **(C-E)** Circadian (C), anti-microbial peptide (D), and *Muc*2 (E) gene expression in DIMS from RC or HF-fed SPF *Reg3γ*^+/−^ or *Reg3γ*^−/−^ mice harvested every 4 hrs over a 12:12 LD. *p<0.05; **p<0.01; ***p<0.001; ****p<0.0001 via Brown-Forsythe and Welch ANOVA followed by Dunnett’s test relative to *Reg3γ*^+/−^ RC control—star color indicates treatment exhibiting significance within ZT time point. ξ indicates significant (p<0.05) co-sinor expression patterns detected via CircWave.

We explored diet vs. REG3γ influences on distal ileum core CC gene expression. Diet or genetic background did not affect overall and diurnal CC expression, further supporting core CC gene network and *Reg3γ* independence (**Figure 4C, S4B; Table S1**). Significantly elevated and diurnal *Reg3γ* expression were evident in RC *Reg3γ*^+/−^ mice, peaking at ZT10, with no detectable transcript in *Reg3γ*^−/−^ mice (**Figure 4D**). Diet or genotype had no effect on levels, diurnal rhythms, or amplitude of *Crypt4*, *Lyz1, Ang4*, and *Tlr* expression (**Figure S4C, Table S1**). RC-fed *Reg3γ*^+/−^ mice exhibited increased overall levels and diurnal patterns of *Muc2* expression, which peaked at ZT10 mirroring *Reg3γ*; however, *Muc2* expression was reduced and arrhythmic in HF-fed *Reg3γ*^+/−^ as well as RC and HF-fed *Reg3γ*^−/−^ mice (**Figure 4E**). These data suggest independence of *Reg3γ* and other AMPs from the core CC, while *Reg3γ* appears to mediate diet effects on *Muc2* expression and rhythmicity, which may impact diurnal rhythms in distal ileum mucosal barrier function.

### HF overwhelms the modest influence of REG3γ on ileal microbiota and shifts specific bacterial families correlated with *Reg3γ* expression

We determined the influence of diet vs. *Reg3γ* status on DILC and DIMS gut microbial community membership and oscillations via 16S rRNA gene amplicon sequencing in samples from *Reg3γ*^+/−^ and *Reg3γ*^−/−^ mice. Regardless of genotype, HF significantly increased relative abundance of Firmicutes with corresponding decrease in Bacteroidetes (**Figure 5A**). Less dominant phyla exhibited modest or no significant change in HF groups (**Figure S5A**). HF also shifted community membership regardless of genotype in both DILC and DIMS assessed via Beta-diversity analyses using Bray Curtis distances (**Figure 5B, S5B**; **Table S3**). This was confirmed via independent comparison of dietary conditions within genotype (**Figure S5C**; DIMS data not shown). Significant differences in DILC community membership were observe between genotypes, but only on RC which indicated a diet-dependent genotype effect (**Figure 5C**, left panel), which was ablated by HF (**Figure 5C**, right panel). No difference in DIMS was observed (data not shown). These data suggest REG3γ exerts a modest but significant influence on community membership under RC, which is overwhelmed by HF.

**Figure 5.**
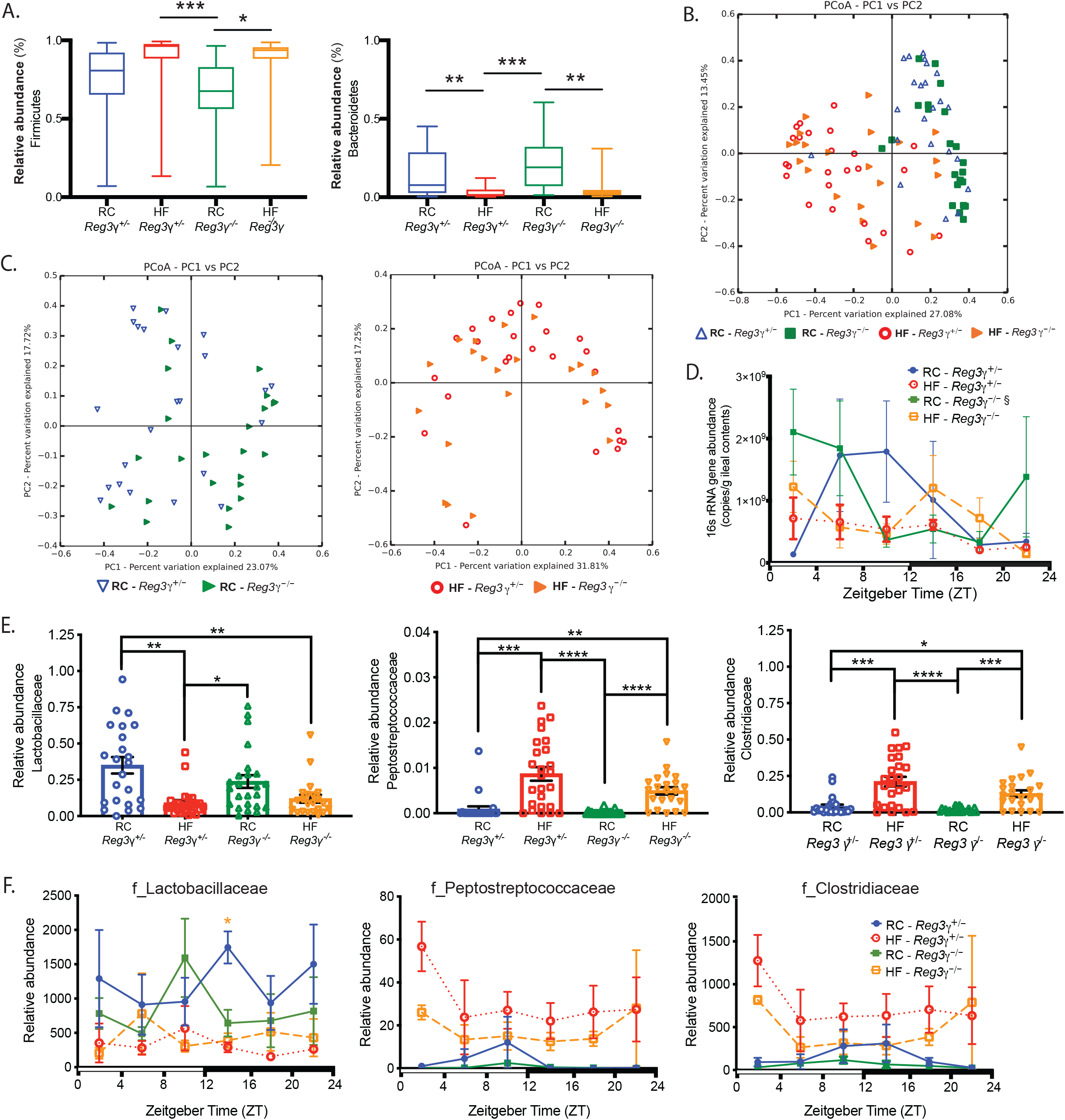
REG3γ deficiency modestly impacts distal SI gut microbe community membership in a diet-dependent manner. **(A)** Relative abundances of dominant bacterial phyla in DIMS from RC or HF-fed SPF *Reg3γ*^+/−^ or *Reg3γ*^−/−^ mice, determined via 16S rRNA gene amplicon sequencing. Box-whisker plots represent mean±min/max. *p<0.05; **p<0.01; ***p<0.001 via Brown-Forsythe and Welch ANOVA followed by Dunnett’s test. **(B,C)** Bray-Curtis PCoA of 16S rRNA amplicon sequencing in DILC from both *Reg3γ*^+/−^ or *Reg3γ*^−/−^ mice fed RC or HF (B) and separated by RC or HF (C). **(D)** 16S rRNA gene copy number in DILC from RC or HF-fed *Reg3γ*^+/−^ or *Reg3γ*^−/−^ mice. Data represent mean±SEM. Statistics via Brown-Forsythe and Welch ANOVA followed by Dunnett’s test relative to *Reg3γ*^+/−^ RC control. ξ indicates significant (p<0.05) co-sinor expression patterns detected via CircWave. **(E)** Average family relative abundances in DILC from RC or HF-fed *Reg3γ*^+/−^ or *Reg3γ*^−/−^ mice. Data represent mean±SEM. *p<0.05; **p<0.01; ***p<0.001; ****p<0.0001 via Brown-Forsythe and Welch ANOVA followed by Dunnett’s test. **(F)** Relative abundance of bacteria families in DILC from RC or HF-fed *Reg3γ*^+/−^ or *Reg3γ*^−/−^ mice. Data represent mean±SEM. *p<0.05 via Brown-Forsythe and Welch ANOVA followed by Dunnett’s test relative to *Reg3γ*^+/−^ RC control—star color indicates treatment exhibiting significance within ZT.

16S rRNA gene copy number in DILC was not different between any group (one-way ANOVA, p>0.05; **Figure S5D**), while diurnal rhythmicity was only apparent in RC-fed *Reg3γ*^−/−^ mice (**Figure 5D, Table S1**). Regardless of genotype, HF significantly altered DILC relative abundances (ANOVA Bonferroni p<0.05, **Table S5**), while diet-dependent differences in oscillation were observed between genotypes (**Figure S2E**). We further explored the impact of diet vs. REG3γ on microbial community membership at the family level. HF resulted in decreased relative abundance of Lactobacillaceae and increased Peptostreptococcaceae and Clostridiaceae regardless of genotype (**Figure 5E**). DIMS were nearly identical, except relative abundance of Clostridiaceae was decreased by HF (**Figure S5F**). DILC Lactobacillaceae relative abundance was decreased in HF-fed *Reg3γ*^−/−^ at ZT14 relative to RC-fed *Reg3γ*^+/−^ mice, while both Peptostreptococcaceae and Clostridiaceae tended to be lower at nearly all ZTs in RC-fed mice regardless of genotype (**Figure 5F**). Although diurnal patterns were similar, Clostridiaceae was less abundant in DIMS at nearly all ZTs (**Figure S5G**). These data confirm that diet broadly reshapes distal ileum microbes with a modest, and diet-dependent effect of REG3γ.

### *Reg3γ* deficiency coupled with HF promotes gain of oscillation in Clostridiales and bacteria that are susceptible to REG3γ’s bactericidal action

Examination of 16S rRNA amplicon sequences from DILC via eJTK revealed RC-fed *Reg3γ*^+/−^ mice displayed significantly more oscillating OTUs than HF-fed counterparts, while both *Reg3γ*^−/−^ groups exhibited nearly identical numbers (**Figure 6A, Table S6**). DIMS exhibited opposite patterns; regardless of diet, *Reg3γ*^−/−^ mice exhibited the greatest oscillating OTUs (**Figure 6B**). Further, RC-fed *Reg3γ*^+/−^ mice exhibited the highest diversity of oscillating DILC OTUs at the taxonomic level of order (**Figure 6C**, left panel). In *Reg3γ*^+/−^ mice, HF induced more oscillating Bacteroidales, fewer Lactobacillales, and no major change in Clostridiales. In *Reg3γ*^−/−^ mice, HF induced fewer oscillating Bacteroidales, more Lactobacillales, and significantly more Clostridiales. DIMS exhibited an inverse microbial oscillator phenotype (**Figure 6C,** right panel). Both RC-fed groups exhibited decreased diversity in taxonomic order of oscillating OTUs and more oscillating Lactobacillales relative to HF (**Table S7**). *Reg3γ*^−/−^ mice also hosted the highest number of oscillating Bacteroidales and Clostridiales relative to *Reg3γ*^+/−^ counterparts.

**Figure 6.**
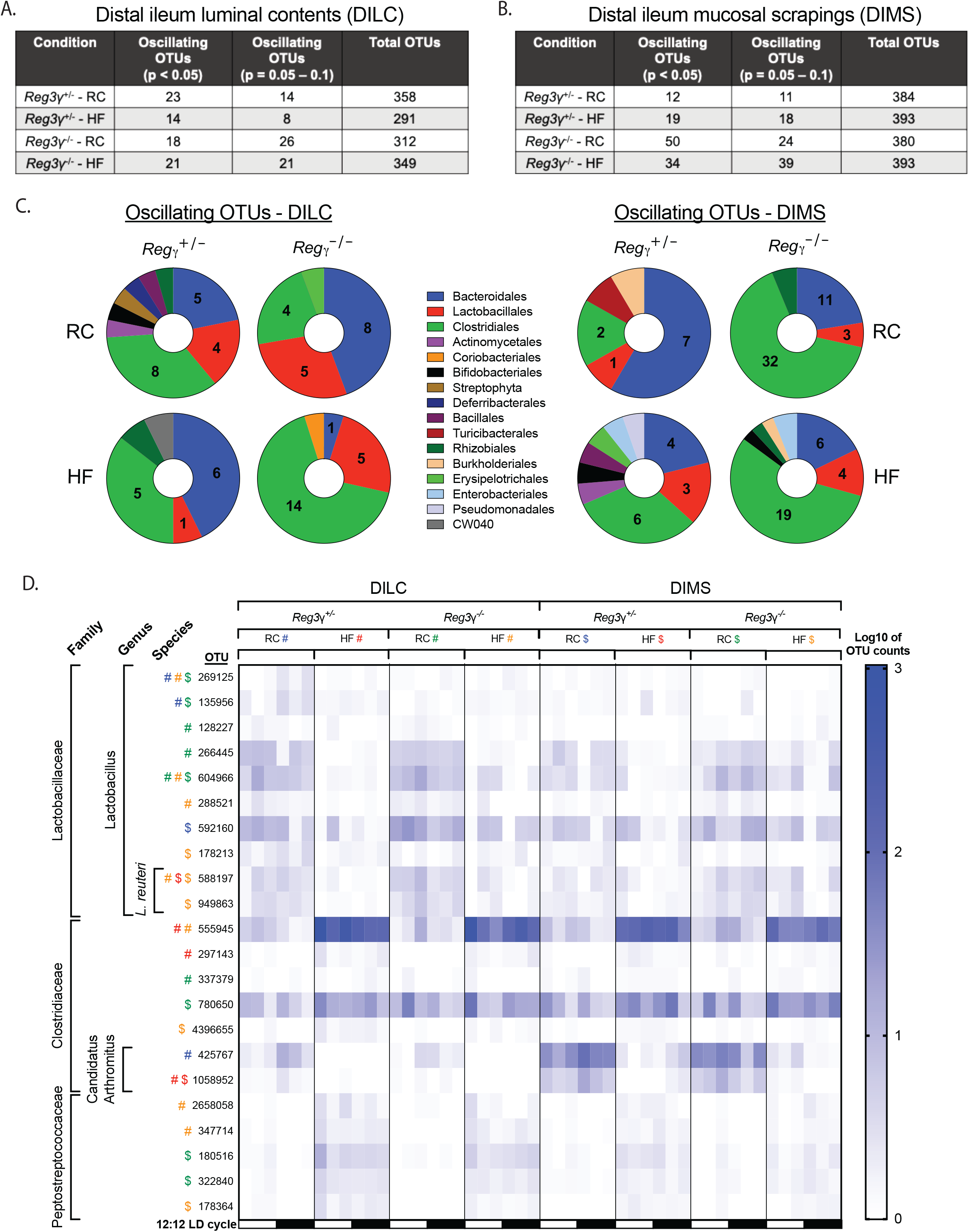
Diet coupled with *Reg3γ* deficiency induces unique microbial community member specific diurnal oscillations. **(A,B)** Number of significantly oscillating OTUs detected via eJTK in DILC (A) and DIMS (B) from RC or HF-fed *Reg3γ*^+/−^ or *Reg3γ*^−/−^ mice. **(C)** Proportion of significantly oscillating OTUs detected via eJTK divided by order in DILC (left) and mucosal scrapings (right) from RC or HF-fed *Reg3γ*^+/−^ or *Reg3γ*^−/−^ mice. Pie charts indicate number of significantly oscillating OTUs. **(D)** Relative abundances of OTUs (Log10 of counts) of bacterial families that exhibit significant oscillations detected via eJTK in DILC and DIMS from RC or HF-fed *Reg3γ*^+/−^ or *Reg3γ*^−/−^ mice. Symbols represent significant oscillations within a group.

We next examined oscillations of Lactobacillaceae, Clostridiaceae, and Peptostreptococcaceae (**Figure 6D**). Many oscillating *Lactobacillus* OTUs exhibited increased relative abundance in RC-fed mice, regardless of genotype. 60% of these OTUs oscillated only in RC-fed mice, indicating *Lactobacillus* diurnal rhythms are dependent on RC. Despite an overall lack of species-level annotation, two *Lactobacillus* OTUs were classified as *L. reuteri*. Although the relative abundance of *L. reuteri* was increased in RC, these OTUs only oscillated in HF when diurnal *Reg3γ* expression was impaired. Similarly, nearly all oscillating Clostridiaceae and Peptostreptococcaceae OTUs were only observed when diurnal *Reg3γ* expression patterns were impaired or completely absent. This expands our findings from SPF WT mice, in which HF only impacted relative abundance patterns and overall number of oscillating microbiota (**Figure 2**). This suggests bacteria that are normally susceptible to REG3γ, i.e., *L. reuteri*, Clostridiaceae, and Peptostreptococcaceae, only gain rhythmicity following HF-diet induced reshaping of distal ileum microbiota coupled with absence of host REG3γ.

## Discussion

Biological rhythms are essential to all life forms for coordination of internal events with the environment to maximize efficiency in metabolic, neural, immune, and other critical functions. CRs consist of a feedback loop of transcription factors that induce rhythmic gene expression patterns to orchestrate a hierarchy of downstream events to achieve this efficiency. However, the gut microbiome, which also exhibits rhythms, cannot entrain to photic environmental cues, instead responding to signals such as meal timing, content, and amount. Three major findings of our study underscore the uniqueness of gut microbial rhythms, as well as provide insight into a dynamic, transkingdom interaction between microbes and local host factors. First, we define a temporal push-pull relationship between specific microbial oscillators and the innate AMP *Reg3γ*, which exhibits diurnal rhythmicity but is independent of the core CC. Second, relevant to human metabolic disorders, we demonstrate HF, low-fiber diet results in loss of functional microbial oscillators and gain of populations that no longer engage with and suppress *Reg3γ*. Third, the combination of HF and loss of *Reg3γ* allows specific microbial oscillators to emerge, which could contribute to metabolic imbalances such as DIO. While an essential role of SI microbes in regulating circadian networks and metabolism has been previously demonstrated, these studies focused on host parameters and outcomes without examining gut microbe membership, function, and rhythmicity. Here, we demonstrate a dynamic, bidirectional interaction between diet-induced gut microbes and *Reg3γ* through the lens of CRs and metabolic homeostasis.

In our model system, we demonstrate the SI core CC gene network exhibits remarkable autonomy from both diet- and microbe-derived cues (**Figure 1A**), confirming previous work (Wang et al., 2017). These results underscore uniqueness of the gut relative to other peripheral tissues; we previously showed rhythmicity of hepatic core CC genes is exquisitely sensitive to diet-induced microbial cues (Leone et al., 2015). Importantly, we show distal SI diurnal *Reg3γ* regulation is instead driven by microbial diurnal cues under RC (**Figure 1B**). HF-induced gut dysbiosis results in arrhythmic *Reg3γ*, underscoring the importance of microbes as key drivers of this phenomenon. The reason for decoupled diurnal regulation of these two systems is unclear; however, it could indicate autonomy of the gut is required to maintain proper timing of essential digestive functions where perturbing rhythmicity of the CC could ultimately compromise host fitness.

Our results clearly demonstrate RC promotes specific microbes that are essential to drive host diurnal SI *Reg3γ* expression. While *Lactobacillus* is known to correlate with *Reg3γ* (Huang et al., 2017), we reveal their diurnal dynamics are intricately intertwined with this phenomenon (**Figure 2F**). In the SI, approximately 15% of microbes oscillate on RC and HF eliminates the majority of oscillators (**Figure 2E**), corroborating previous findings in the distal GI tract (Leone et al., 2015; Thaiss et al., 2014; Zarrinpar et al., 2014). We reveal DIO is not only associated with decreased SI *Reg3γ* expression (Everard et al., 2013, 2014; Vaishnava et al., 2011), but also HF-induced gut dysbiosis is essential to suppress rhythmic *Reg3γ* expression (**Figure 1B**). Our work supports diet overwhelms genetics in shaping microbial SI membership, where HF elicits greater influence than *Reg3γ* genetic deletion (**Figures 2, 5, 6**), confirming previous studies in the distal GI tract (Carmody et al., 2015; Devkota et al., 2012). Interestingly, HF coupled with global REG3γ deficiency permits diet-induced gut microbes to increase proportionally and gain oscillations within the community (**Figure 6D**). Together, this reveals diet is the main driver of gut microbiota membership and oscillations, while *Reg3γ* serves a secondary role in shaping rhythmicity of key SI gut microbes.

We provide direct evidence that diet-induced representative and indigenous SI bacteria associate with and directly induce diurnal *Reg3γ* expression requiring MyD88 (Natividad et al., 2013; Vaishnava et al., 2008). While *Lactobacillu*s correlate with diurnal *Reg3γ* expression, the precise factor they secrete or produce *in vivo* to drive this is not clear. However, we show small molecules derived only from specific *Lactobacillu*s can induce *Reg3γ in vitro* (**Figure S3C**), and may involve direct activation of IEC TLRs. Whether *Lactobacillus*-mediated diurnal *Reg3γ* expression also requires interleukin-22/STAT3-mediated induction is unclear and requires further exploration. Functionally, previous work showed bacteria expressing Bile Salt Hydrolase (BSH), such as *Lactobacillus*, are associated with *Reg3γ* expression (Joyce et al., 2014; O’Flaherty et al., 2018). Whether BSH mediates production of small molecules that directly induce *Reg3γ* expression is unknown. However, the ability of *Lactobacillus* species to express BSH could influence their diurnal oscillations, which may relate to food intake timing and bile acid delivery/reuptake in the ileum (Zarrinpar et al., 2014).

To our knowledge, negative associations between SI *Reg3γ* expression and specific gut microbes have not been well established. We show Clostridiaceae and Peptostreptococcaceae family members do not induce, and may even suppress *Reg3γ* expression *in vitro* in certain instances (**Figures 3G,H**). Whether this occurs *in vivo* remains to be determined. We were unable to successfully monoassociate RC or HF-fed GF mice with *P. stomatis* despite several attempts. Since *P. stomatis* was isolated from human oral cavity, it is possible that murine gut is not a suitable environment and indigenous Peptostreptococcaceae strains could be used in future studies to explore their influence on host *Reg3γ*.

While REG3γ’s ability to target commensal vs. pathogenic bacteria has been shown (Cash et al., 2006a), we also reveal its antimicrobial action may be more targeted. While some *Lactobacillus* species evade REG3γ, *L. reuteri* is quite susceptible (**Figure 3I-K**), more closely aligning with *P. stomatis* and *E. faecalis* (**Figure 3G,H**). This is in line with work showing *Reg3γ* overexpression in murine IECs led to expansion of *Lactobacillus* abundance in general, while certain species such as *L. reuteri*, were reduced (Huang et al., 2017). The reason for *Lactobacillus* species differential sensitivity is unclear, however we posit diurnal induction of and resistance to REG3γ may aid in niche maintenance within the community while contributing to host intestinal health. Given the heterotrophic nature and heterogeneous proteolytic capacity of *Lactobacillus*, it is possible certain species are able to utilize host-derived peptides, including AMPs, as a novel fuel source under nutrient-limiting conditions.

The dual function of REG3γ as both an AMP and hormonal signal remains poorly defined, particularly in HF DIO. We show that global *Reg3γ* deficiency worsens glucose sensitivity under RC feeding conditions and is not exacerbated by HF (**Figure 4B**), which is unique relative to a previous report (Bluemel et al., 2018). While compelling, we cannot establish which tissue compartment drives glucose tolerance and whether diurnal *Reg3γ* expression is important since we employed a global knockout. Further investigation is required to elucidate the direct diurnal signaling influence of *Reg3γ*, locally and globally, and its impact on metabolism. Further, we identified modest differences in luminal SI microbes between RC-fed *Reg3γ*^+/−^ and *Reg3γ*^−/−^ mice, while others have not observed this (Vaishnava et al., 2011). This could be due to microbiome differences between animal vivaria, housing conditions (individual vs. group), our use of heterozygote mice as controls.

Together, we show diet coupled with REG3γ, through their actions on gut microbial populations, are essential components for regulating rhythmicity of specific bacteria, independent of archetypal CC gene networks. These microbial rhythms depend on dynamic diet-host-microbe interactions. We speculate specific bacteria either flourish or are lost as a result of altered dietary intake leading to *Reg3γ* induction or suppression at crucial times. If improper timing or loss of diurnal *Reg3γ* persists, it could render the host more susceptible to infection or worse pathogenesis of chronic diseases, such as DIO. These interactions are a prime example of transkingdom co-evolution that is essential for the mammalian host to adapt to changes in daily dietary intake. The knowledge gained here provides a framework for identification of interventions that restore diurnal host-microbe interactions to alleviate metabolic diseases associated with Western diet.

## Acknowledgements

The present research was supported by NIH NIDDK K01 DK111785 (VL); NIDDK Digestive Diseases Research Core Center (NIH P30 DK42086); NIDDK F31 DK122714 (KF) and the University of Chicago GI Research Foundation. We thank the UChicago Human Tissue Resource Center for histological processing and Gnotobiotic Research Animal Facility staff for animal husbandry. We are indebted to Dr. Lora Hooper for thoughtful discussions regarding experiments and data.

## Author Contributions

KF, EZ, EBC, and VL conceptualized experiments. AK, EZ, JFP, SM, JM, DR, DH, KY, CC, MWM, and VL performed experiments. VL, AK, EZ, KF, and NH analyzed data. KF, EBC, and VL wrote the manuscript. VL oversaw the entire project.

## Supplemental Tables and Figure Titles and legends

**Figure S1, related to Figure 1.**
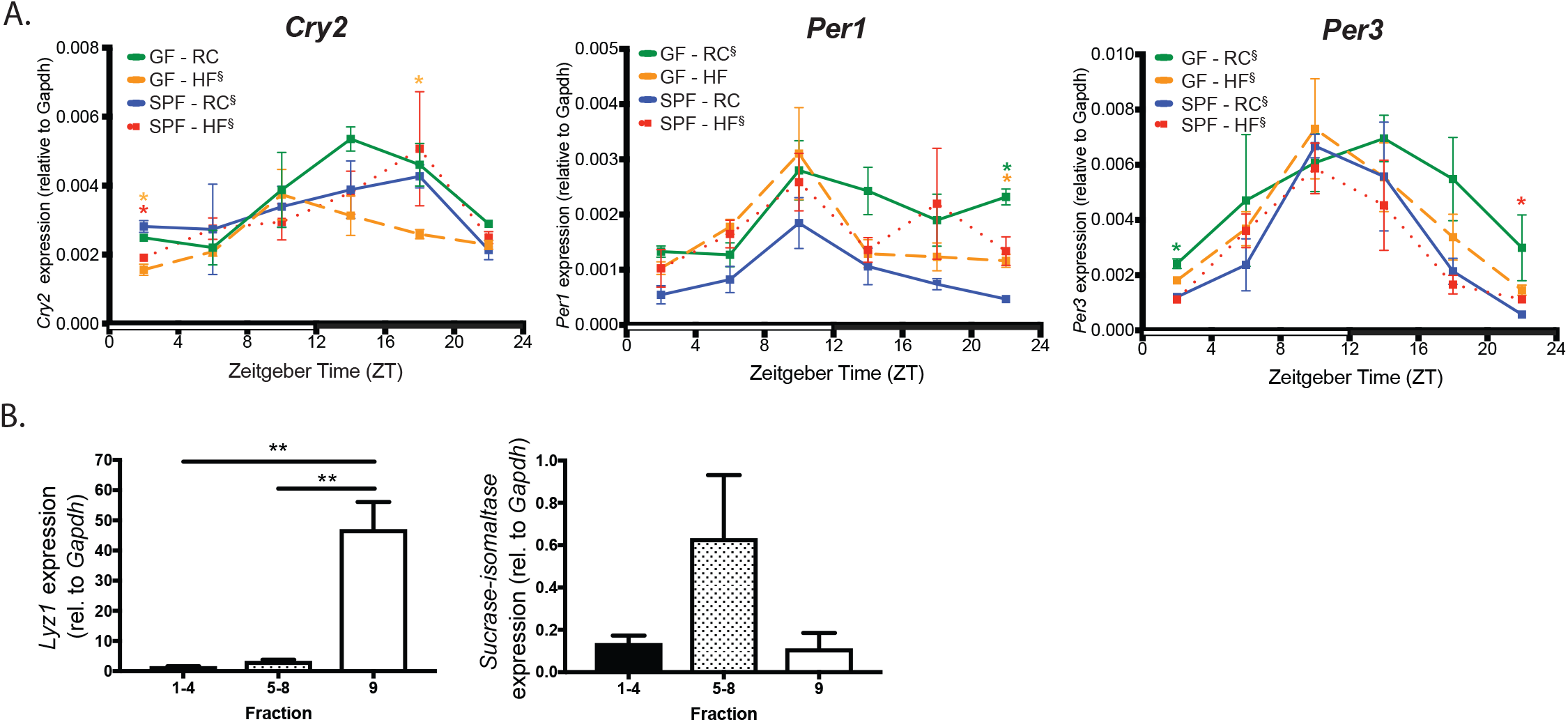
Core CC gene expression is not influenced by diet or gut microbes. **(A)** Diurnal circadian gene expression from DIMS collected every 4hrs from GF and SPF mice fed RC or HF. *p<0.05, **p<0.01, ***p<0.001 via Brown-Forsythe and Welch ANOVA followed by Dunnett’s test as compared to SPF-RC control—star color indicates treatment exhibiting significance within ZT time point. ^ξ^in figure legends indicate significant (p<0.05) co-sinor expression patterns detected via CircWave. **(B)** *Lyz1* and *Sucrase-Isomaltase* expression relative to *Gapdh* in distal ileum epithelial fractions from SPF RC or HF-fed mice. Fractions 1-4 and 5-8 contain absorptive enterocytes, fraction 9 contains intestinal crypts harboring stem cells and Paneth cells. **p<0.01 via Brown-Forsythe and Welch ANOVA followed by Dunnett’s test.

**Figure S2, related to Figure 2.**
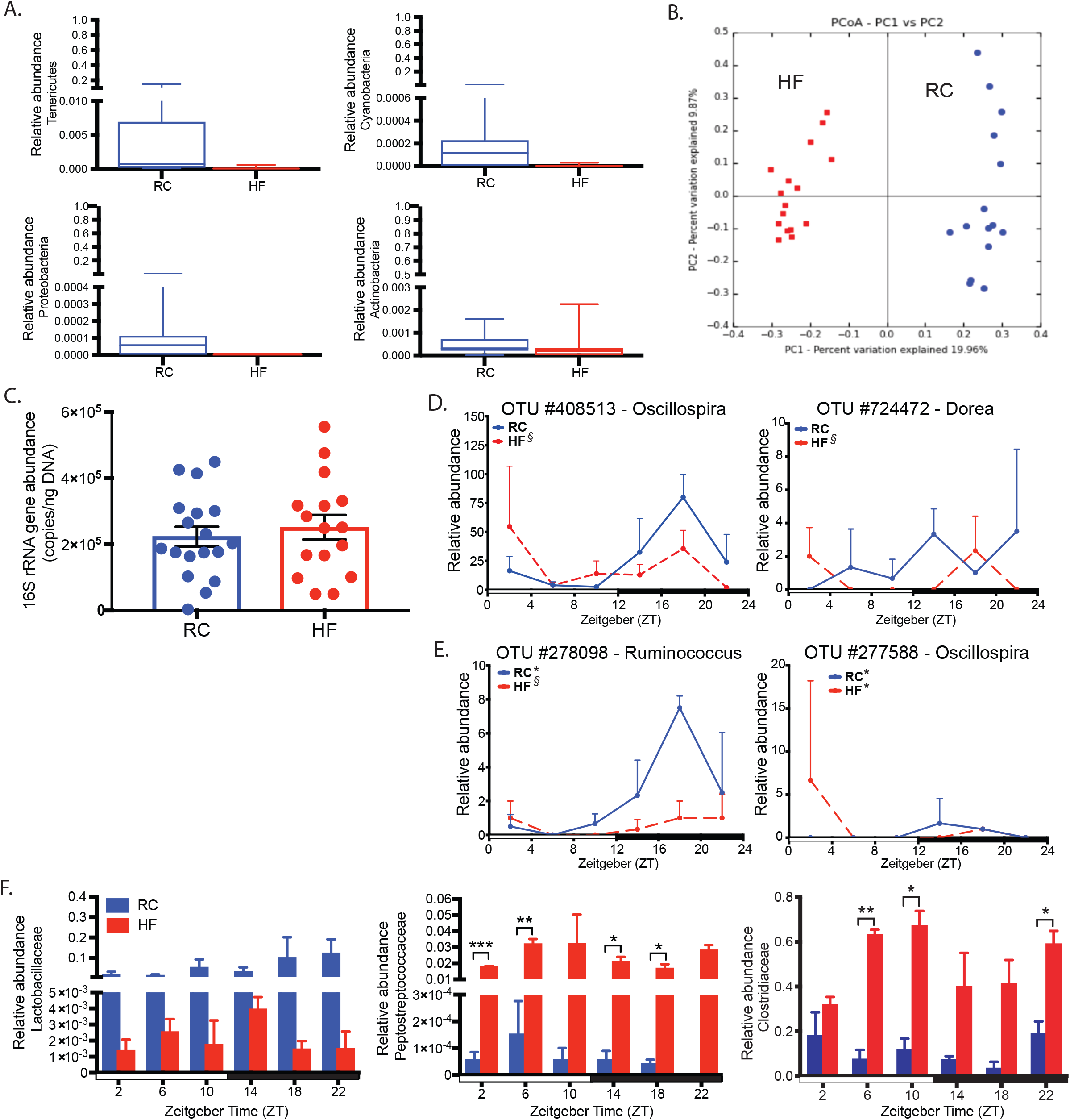
Diet drives differential community membership in non-dominant phyla and at the family level across all time points. **(A)** Relative abundance of less dominant bacterial phyla of DILC from RC or HF-fed SPF mice determined via 16S rRNA amplicon sequencing. Box-whisker plots represent mean ± min/max. Statistics via Welch’s t test. **(B)** PCoA of Canberra distances from 16S rRNA amplicon sequences indicated by diet in DILC. **(C)**16S rRNA gene copy number in DILC from RC or HF-fed mice. Data represent mean ± SEM. n.s.=not significant via Welch’s t test. **(D)** Relative abundances of OTUs mapped to the level of genus that significantly oscillate only in DILC of HF-fed mice. Relative abundances of these OTUs are also shown for RC-fed mice as a comparison. Data represent mean ± SEM. * indicates significant oscillation (p<0.05), while ξ indicates significant oscillation (p=0.05 − 0.1) detected via eJTK. **(E)** Relative abundances of OTUs mapped to the level of genus that significantly oscillate in DILC of RC and HF-fed mice. Data represent mean ± SEM. * indicates significant oscillation (p<0.05), while ξ indicates significant oscillation (p=0.05 − 0.1) detected via eJTK. **(F)** Relative abundance of indicated bacterial family in DILC from RC and HF-fed mice that include OTUs that positively or negatively correlate with *Reg3γ* expression. Data represent mean ± SEM. *p<0.05; **p<0.01; ***p<0.001 via Welch’s t test.

**Figure S3, related to Figure 3.**
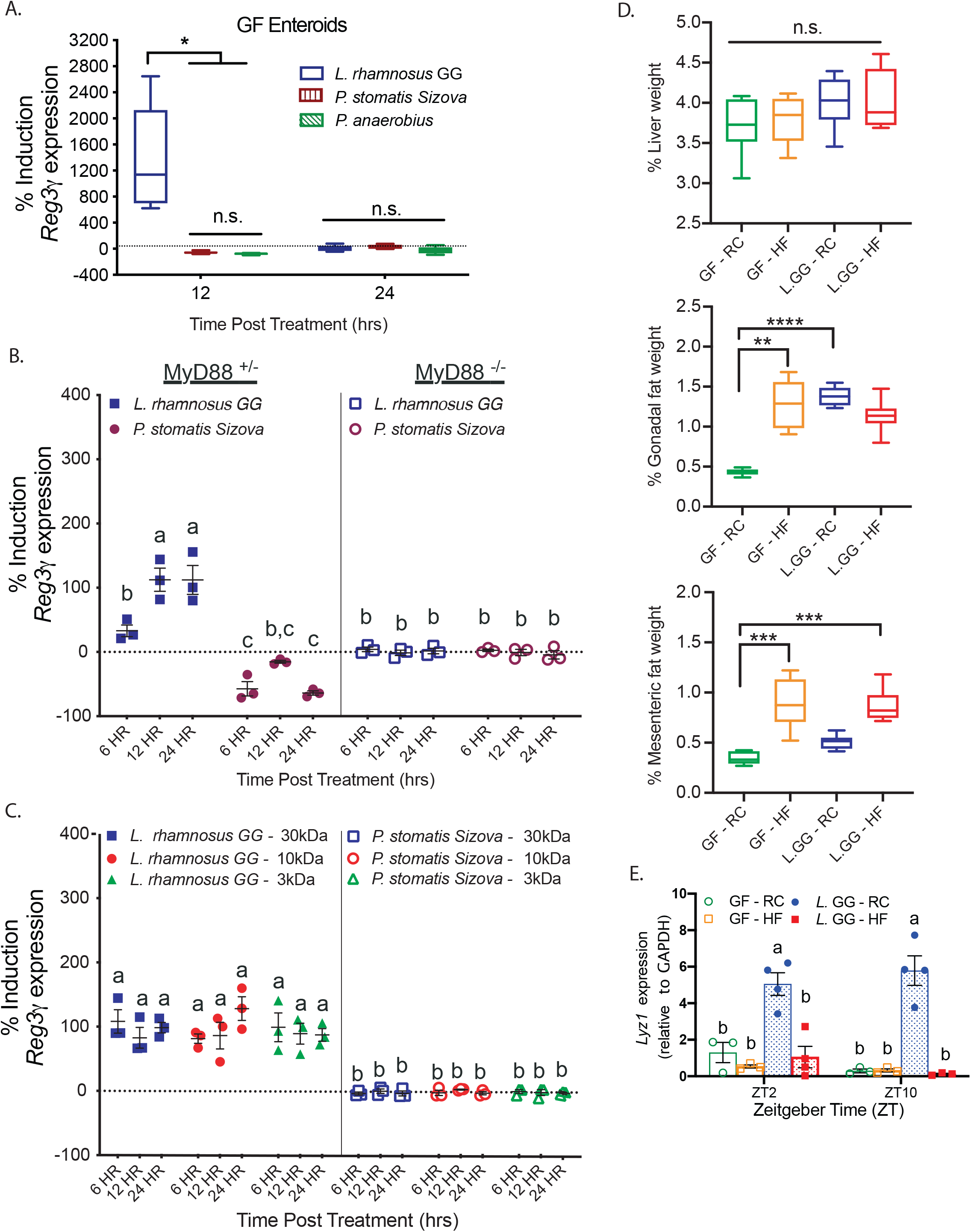
Gram-positive bacteria drive host *Reg3γ* expression in a MyD88-dependent manner *in vitro* and *in vivo*. **(A)** Percent induction of *Reg3γ* expression in enteroids derived from RC-fed GF WT mice following 12 or 24-hr exposure to bacterial lysate from cultured strains LGG, *P. stomatis*, or *P. anaerobius* relative to their respective blank media controls. *n*=3-6 technical replicates/treatment, representative of 3 independent experiments. Box-whisker plots represent mean±min/max. *p<0.05 via Brown-Forsythe and Welch ANOVA followed by Dunnett’s test; n.s.=not significant. **(B)** Percent induction of *Reg3γ* expression of enteroids derived from SPF MyD88^+/−^ or MyD88^−/−^ RC-fed mice. Enteroids were treated with CM from LGG or *P. stomatis* for 6, 12, or 24hrs. *n*=3 technical replicates/treatment. Data presented as mean ± SEM, representative of 2 independent experiments. Columns with the same letter are not significantly different (p>0.05) as determined via two-way ANOVA followed by Tukey’s test. **(C)** Percent induction of *Reg3γ* expression in enteroids derived from RC-fed SPF WT mice following exposure to size fractionated CM from LGG or *P. stomatis* for 6, 12, or 24hrs. *n*=3 technical replicates/treatment. Data are presented as mean ± SEM and are representative of 2 independent experiments. Columns with the same letter are not significantly different (p>0.05) via two-way ANOVA followed by Tukey’s test. **(D)** Liver, gonadal fat pad, and mesenteric fat pad weight expressed as a percent of body weight of GF or LGG-monoassociated mice fed RC or HF. Box-whisker plots represent mean±min/max. *n*=6-8 mice/treatment. *p<0.05, ***p<0.001, ****p<0.0001 via Brown-Forsythe and Welch ANOVA followed by Dunnett’s test. **(E)** *Lyz1* expression in DIMS of GF or LGG-monoassociated WT mice fed RC or HF. Data presented by mean ± SEM, representative of 2 independent experiments. Bars with the same letter are not significantly different (p>0.05) via two-way ANOVA followed by Tukey’s test.

**Figure S4, related to Figure 4.**
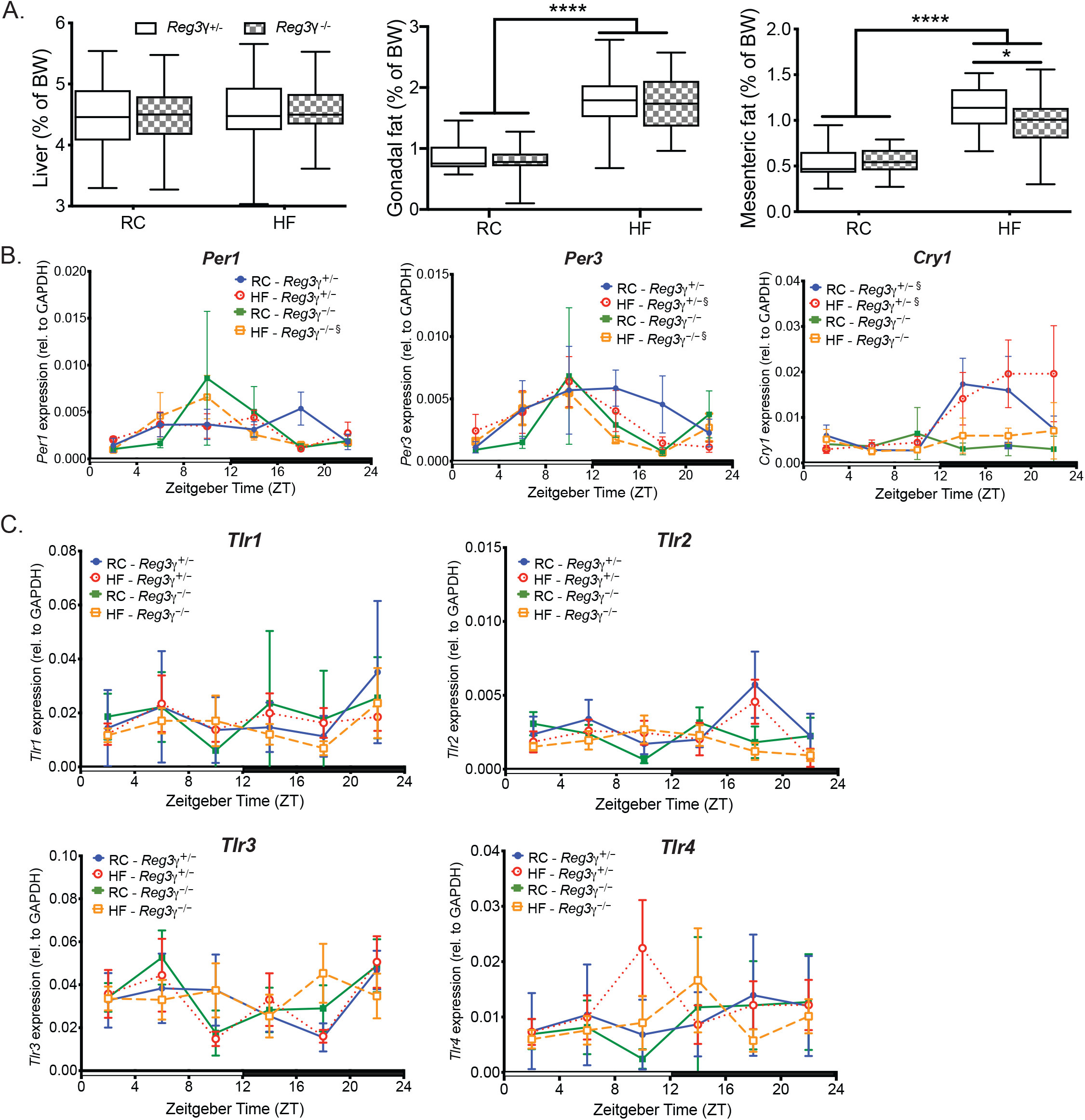
*Reg3γ* deficient mice are equally susceptible to HF diet-induced body composition changes with no changes in core CC or TLR expression. **(A)** Liver, gonadal fat pad, and mesenteric fat pad weight expressed as a percent of body weight of *Reg3γ*^+/−^ or *Reg3γ*^−/−^ mice fed RC or HF. Box-whisker plots represent mean±min/max. *n=*24-25 mice/treatment. *p<0.05; ****p<0.0001 via two-way ANOVA followed by Tukey’s test. **(B-C)** Diurnal circadian (B) and TLR (C) gene expression in DIMS from *Reg3γ*^+/−^ or *Reg3γ*^−/−^ mice fed RC or HF. Statistics via Brown-Forsythe and Welch ANOVA followed by Dunnett’s test relative to *Reg3γ*^+/−^ RC control—star color indicates treatment exhibiting significance within timepoint. ξ indicate significant (p<0.05) co-sinor expression patterns detected via CircWave.

**Figure S5, related to Figure 5.**
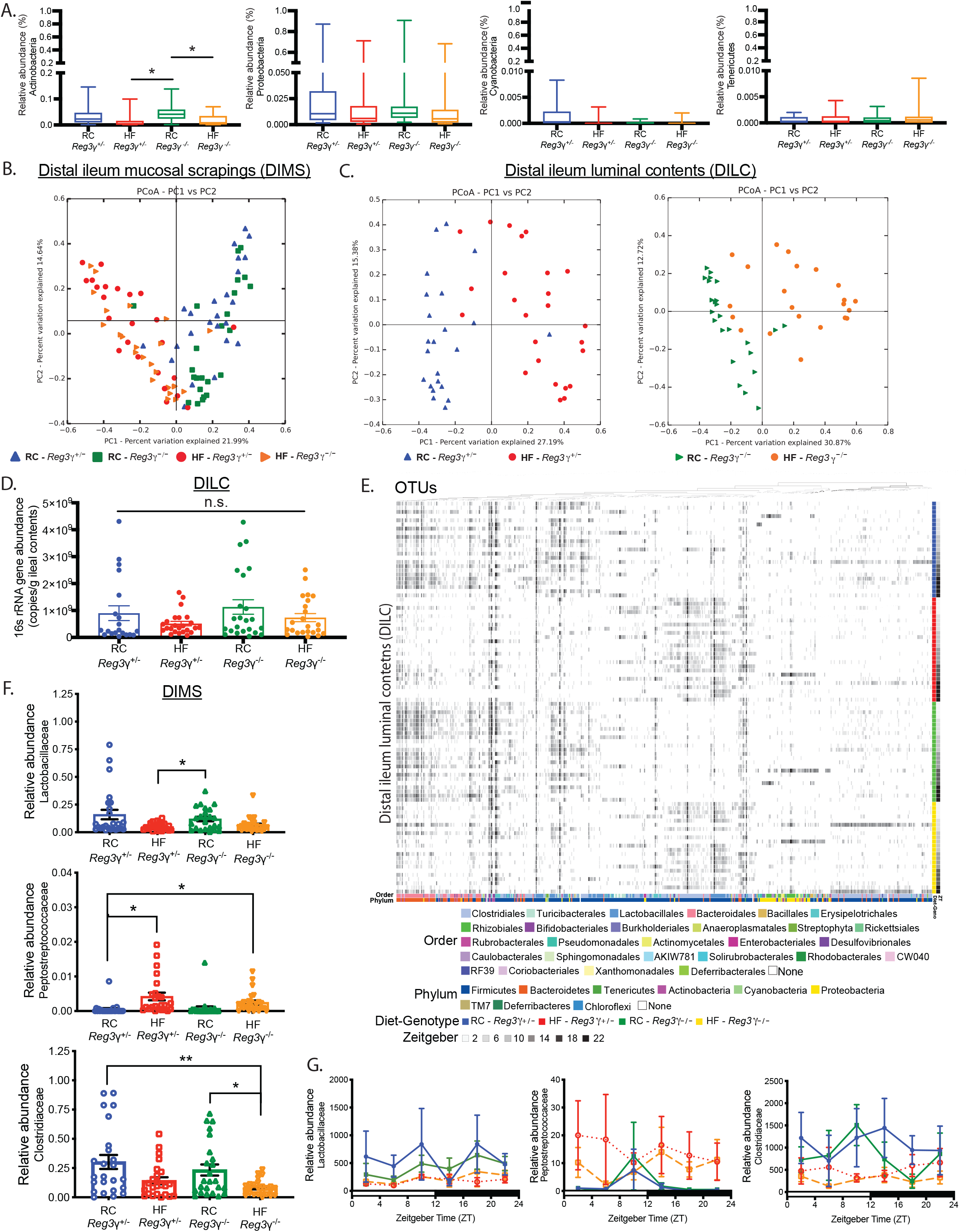
Diet is the primary driver of overall microbial community membership regardless of host *Reg3γ* status. **(A)** Relative abundances of less dominant bacterial phyla of DILC from RC or HF-fed *Reg3γ*^+/−^ or *Reg3γ*^−/−^ mice via 16S rRNA amplicon sequencing. Box-whisker plots represent mean±min/max. *p<0.05 via Brown-Forsythe and Welch ANOVA followed by Dunnett’s test. **(B)** Bray-Curtis PCoA of 16S rRNA amplicon sequencing of DIMS from *Reg3γ*^+/−^ or *Reg3γ*^−/−^ mice fed RC or HF. **(C)** Bray-Curtis PCoA of 16S rRNA amplicon sequencing of DILC from RC- and HF-fed mice, separated by *Reg3γ*^+/−^ or *Reg3γ*^−/−^. **(D)** 16S rRNA gene copy number in DILC from *Reg3γ*^+/−^ or *Reg3γ*^−/−^ mice fed RC of HF. Data represent mean ± SEM. n.s.=not significant via Brown-Forsythe and Welch ANOVA followed by Dunnett’s test. **(E)** OTU 16S rRNA relative abundances in DILC from *Reg3γ*^+/−^ or *Reg3γ*^−/−^ mice fed RC or HF (*n*=3-4 mice/timepoint) using anvi’o. Columns represent OTUs, rows represent samples. Colored bars at bottom represent taxonomic assignment. Colored bars at right represent diet, gray bars represent ZT. **(F)** Average relative abundance of indicated bacterial family in DIMS from *Reg3γ*^+/−^ or *Reg3γ*^−/−^ mice fed RC or HF. Data represent mean ± SEM. *p<0.05; **p<0.01 via Brown-Forsythe and Welch ANOVA followed by Dunnett’s test. **(G)** Relative abundance of indicated bacterial family in DIMS from *Reg3γ*^+/−^ or *Reg3γ*^−/−^ mice fed RC or HF. Data represent mean ± SEM. Statistics via Brown-Forsythe and Welch ANOVA followed by Dunnett’s test relative to *Reg3γ*^+/−^ RC control.

## Experimental procedures (STAR Methods)

### Lead contact

Further information and requests for resources and reagents should be directed to and will be fulfilled by the Lead Contact, Vanessa Leone.

### Materials Availability

Frozen stocks of *Lactobacillus* species isolated and cultivated in the present study are stored in the facility of the Lead Contact.

## Experimental Model and Subject Details

### Mice

All animal protocols and experimental procedures were approved by the University of Chicago Institutional Animal Care and Use Committee (IACUC). Specific pathogen-free (SPF) C57Bl/6 male mice, aged 8-10 weeks, were purchased from Jackson Laboratories (Bar Harbor, ME, USA). Male, age-matched SPF *MyD88*^+/−^, *MyD88*^−/−^, *Reg3γ*^+/−^ and *Reg3γ*^−/−^ as well as germ-free (GF) WT mice on a C57Bl/6 background were bred in the University of Chicago animal vivarium. GF mice were maintained in plastic flexible film isolators (CBC Ltd. Madison, WI, USA). All mice were held under standard 12:12 light/dark conditions (lights on beginning at 6am, Zeitgeber (ZT) 0) and individually housed. After two weeks acclimatization, mice were allotted to one of two dietary treatments: regular, low-fat (RC) chow (10% dietary fat, TD.2018S, Envigo, Madison, WI, USA) or high fat (HF) diet (37.4% dietary fat, TD.97222, custom diet, Envigo, Madison, WI, USA) (see supplemental information in Leone et al., 2015 for detailed dietary components). GF diets were irradiated and tested before and after experiments for sterility. Macronutrient distribution of experimental diets is listed in Table S11. All mice were fed *ad libitum*. Body weights and food consumption were monitored weekly throughout the study. After 4 weeks, mice were sacrificed via CO_2_ asphyxiation followed by cervical dislocation over 24hrs at six ZT time points: ZT 2=8 AM, ZT 6=12 PM, ZT 10=4 PM, ZT 14=8 PM, ZT 18=12 AM, and ZT 22=4 AM. DILC and DIMS were snap frozen in liquid nitrogen and stored at −80C until further analyses.

### Preparation of isolated epithelial cells

A modified version of the Weiser method was utilized to isolate epithelium from the villus-crypt axis (Weiser, 1973). Briefly, the distal 11cm of the ileum was isolated from RC or HF-fed mice at ZT2 or 10. Sections were perfused with ice cold PBS plus 1mM DTT, followed by eversion, tied at one end, and filled to distension with ice cold PBS. Ileal segments were then incubated at 37C for 15min in 15mL citrate buffer , followed by transfer to a PBS buffer containing 1.5mM EDTA, 0.5mM DTT, and 1mg/mL bovine serum albumen and shaken at 175 RPM at 37°C for 10 min. Intestinal segments were then transferred to fresh PBS buffer solution and incubated nine consecutive times for 10, 6, 5, 5, 9, 10, 15, 25, and 30 min., respectively as previously described (Ferraris et al., 1992). Fractions 1-4 and 5-8 were pooled and considered villi while fraction 9 was considered crypt, containing Paneth cells. Combined fractions were spun for one minute at 13,000G and resuspended in Trizol, followed by RNA extraction and cDNA synthesis for qPCR analysis as described below.

### Enteroid culture

Enteroids were grown as previously described (Sato et al., 2011). Briefly, the distal 11cm of the SI was removed from GF or SPF C57Bl/6 WT, *MyD88*^+/−^, or *MyD88*^−/−^ mice and opened longitudinally. Tissues were rinsed in ice-cold PBS to remove contents. A glass slide was used to gently remove the villi, and the mucosa was minced into 1- to 2-mm pieces with a scalpel and collected into 10ml of ice-cold PBS. Pieces were agitated and rinsed several times using a serum-coated serological pipette. Intestinal pieces were resuspended in 25ml of ice-cold 2.5 mM EDTA-PBS and rotated at 4°C for 30min. Following incubation, EDTA-PBS was replaced with 10 ml of Advanced DMEM/F12 (ADF) medium (Thermo Scientific). After gently disrupting with a pipette three times, supernatant was discarded and fresh ADF medium was added and repeated three times. Cells were centrifuged at 300 *g* at 4°C for 5 min, resuspended in 10ml ADF medium, and passed through a 70-μm cell strainer to remove debris. Cells were centrifuged at 300 *g* at 4°C for 3 min and resuspended in complete ADF medium containing GlutaMAX (Thermo Scientific), HEPES buffer (Thermo Scientific), penicillin-streptomycin (Thermo Scientific), N2 supplement (Thermo Scientific), B-27 Supplement Minus Vitamin A (Thermo Scientific), murine EGF (50 ng/ml; Thermo Scientific), noggin (100 ng/ml; Peprotech, Rocky Hill, NJ), jagged-1 (1 μM; Anaspec), Y27632 (10 nM; Cayman Scientific, Ann Arbor, MI, USA), and R-spondin-1 (500 ng/ml; Peprotech). The cell pellet was resuspended in a ratio of 1:2 ADF to Matrigel (BD Biosciences, San Jose, CA, USA) and plated onto a pre-warmed, collagen-coated, 24-well-plate. Matrigel beads were allowed to solidify for 1 hr at 5% CO_2_ at 37°C before adding 500 μl ADF culture media.

### Bacterial culture

*E. faecalis, E. coli K12, L. rhamnosus GG, L. reuteri, L. intestinalis, L. murinus*, and *L. johnsonii*. *P. stomatis* Sizova, *P. anaerobius*, *B. thetaiotamicron*, and were streaked out from frozen glycerol stocks onto agar-containing plates of either brain heart infusion media (BHI, *E. faecalis)*, Luria-Bertani media (LB*, E.coli* K12), BHI-Supplemented media (BHIS, *P. stomatis* Sizova), RCM media (*P. anaerobius*), or de Man, Rogosa and Sharpe media (MRS, *Lactobacillus* strains). Single colonies were inoculated into their respective liquid media and cultivated under aerobic or anaerobic conditions (Coy Laboratory Products, Inc., Grass Lake, MI) at 37°C for 24-48hrs. Genomic DNA of indigenous strains *L. reuteri, L. intestinalis, L. murinus*, and *L. johnsonii* were isolated from individual colonies. Forward and reverse 16S rRNA gene sequences were obtained using universal PCR primers (8F and 1492R), joined, and BLAST results were used to identify most closely related species. See Key Resources Table for detailed information on each strain.

## Method Details

### Quantitative PCR

Total RNA was isolated by homogenizing tissue using TRIzol (1559018, Ambion, Hampton, NH) and chloroform extraction method, as previously described (Leone et al., 2015). RNA purity was validated through UV-Vis spectrophotometry via Nanodrop Lite (Thermo Scientific, Wilmington, DE, USA). 1μg of total RNA was reverse-transcribed to complementary DNA (cDNA) using the Transcriptor First Strand cDNA Synthesis Kit (Roche, Indianapolis, IN, USA) according to manufacturer’s instructions. Relative quantification of gene expression was performed using a LightCycler 480 Real-Time PCR System (Roche). Forward and reverse primers were combined with SYBR Green PCR Supermix (Bio-rad) and nuclease-free water to amplify the following host-specific gene expression: *Reg3γ, Lysozyme1*, *Cryptidin4*, *Ang4*, *Tlr1, 2, 3, 4, 9, Clock*, *Bmal1*, *Per1-3*, *Cry1-2, Muc2*, *Sucrase Isomaltase*, and *GAPDH* (see Key Resources Table for primer sequences). Gene expression data are presented as 2^−ΔCt^ (housekeeping gene – target gene), where the house-keeping gene is GAPDH.

### Western Blot

To prepare protein for Western blot, 5 mg of tissue from distal ileal mucosal scrapings was lysed in 250 μL of ice-cold protein lysis buffer (Cell Signaling Technology Cell Lysis Buffer, cOmplete Mini Protease Inhibitor Cocktail, Sigma-Aldrich, 100μM PMSF). 20-30μg of protein extract was separated on a 4-20% precast polyacrylamide gel and transferred to a PVDF membrane (Millipore). Membranes were blocked with 5% nonfat milk in Tris-buffered saline (TBS) (20 mM Tris pH7.6, 150 mM NaCl) and incubated overnight at 4°C in 2% nonfat milk in Tris-buffered saline-Tween (TBS-T) (20 mM Tris pH7.6, 150 mM NaCl, 0.1% Tween-20) containing primary antibodies: anti-REG3γ (1:500; Abcam, Cambridge, MA), anti-GAPDH (1:1000; Invitrogen). Membranes were washed three times for 5 min in TBS-T and incubated for 1 hr at room temperature in 2% nonfat milk in TBS-T containing goat anti rabbit Alexa Fluor 680, Donkey anti-mouse Alexa Fluor 790 secondary antibody (1:100,000 Abcam). Membranes were washed three times for 5 min each in TBS-T and imaged using the LI-COR Odyssey (LI-COR Biosceinces).

### Tissue Histology and Immunofluorescence

For immunofluorescence, sections were fixed in 4% paraformalydehyde overnight. Fixed tissue sections were processed (Tissue-Tek VIP, Sakura Finetek, Torrance, CA) and embedded in paraffin. Five μm sections were cut and mounted on charged glass slides, and deparaffinized. To visually localize and semi-quantitatively measure REG3γ protein within the ileal epithelium and Paneth cells, immunofluorescence was performed on paraformalydehyde fixed sections. Following deparaffinization and rehydration, antigen retrieval was performed by boiling slides in a 10 mM/L sodium citrate bath (pH 6.0). Slides were then blocked with 10% bovine serum albumin (BSA)-PBS for 1 hr. Samples were incubated with primary antibody for Lysozyme (1:400, ab108508, rat monoclonal IgG, Abcam) or REG3γ (1:500, ab198216, rabbit polyclonal IgG, Abcam) overnight in 1% BSA-PBS at 4°C in a humidified chamber. Remaining solutions were washed and samples were incubated with respective secondary antibodies in 1% BSA-PBS (1:1000, Alexa Fluor 594; Invitrogen, Grand Island, NY). Slides were imaged following DAPI staining and coverslip placement.

### 16S DNA Extraction, Sequencing, and Analysis

Distal ileum luminal contents (DILC), and ileal mucosal scrapings (DIMS) were collected in screw cap tubes as previously described in DNA lysis buffer (Leone et al., 2015). After addition of 0.1-mm-diameter zirconia/silica beads (BioSpec Products, Bartlesville, OK, USA), samples were disrupted using a Mini-Beadbeater-8k Cell Disrupter (BioSpec Products). Supernatants were extracted with an equal volume of Phenol:Chloroform:Isoamylalcohol (25:24:1; Ambion, Austin, TX, USA) and DNA precipitated using an equal volume of 100% ethanol. DNA concentration was then measured via Nanodrop Lite (Thermo Scientific, Wilmington, DE, USA) and subsequently diluted to 25 μg/μl. The V4-V5 region of the 16S rRNA encoding gene was amplified using standard Earth Microbiome Project protocols. Sequencing was performed at the High-Throughput Genome Analysis Core (HGAC; part of the Institute for Genomics & Systems Biology [IGSB]) at Argonne National Laboratory. Forward and reverse reads were joined, trimmed and aligned using EA-utils, then classified using the Quantitative Insights Into Microbial Ecology (QIIME) toolkit (Caporaso et al., 2010). OTUs were picked at 97% sequence identity using open reference OTU picking protocol against the Greengenes database. These representative sequences were aligned using PyNAST and taxonomy was assigned using the RDP Classifier. The PyNAST-aligned sequences were also used to build a phylogenetic tree with FastTree and Bray Curtis and Canberra distances were used to statistically compare beta-diversity, and visual comparisons were performed via Principal Coordinate Analysis (PCoA) ordination. To evaluate which microbial taxa exhibited rhythmic oscillations, empirical-JTK cycle with asymmetry (eJTK) software was implemented (Hutchison et al., 2015). Spearman and Pearson correlation was performed to determine which microbial taxa were correlated with host *Reg3γ* gene expression. Analysis and Visualization Platform for ‘Omics data (anvi’o) (Eren et al., 2015) was used to create at heatmap for visually assessing 16S rRNA gene changes in taxonomic membership and abundance by diet and ZT.

### Bacterial Gene Quantification

16S rRNA gene copy number was determined from DILC and DIMS as previously described (Leone et al., 2015). Genes were quantified by determining a standard curve for gene copy number by cloning 16S sequence into pCR4-TOPO plasmid (see Key Resources Table for primer sequences).

### Enteroid stimulation with bacterial culture conditioned media

Stimulation of enteroids with bacterial culture media was performed by adding 10% of filter-sterilized conditioned media (CM) collected from log-phase *Lactobacillus rhamnosus GG* (ATCC 53103), *P. stomatis* (ATCC BAA-2664 CM2), or *P. anaerobius* to complete ADF enteroid culture media. CM treatment was compared to blank sterile-filtered media controls. Treated enteroid culture plates were kept at 5% CO_2_ at 37°C until collection at 6, 12, and 24hrs post-treatment. RNA collection, cDNA synthesis, and qPCR analysis were performed as described above.

### Monoassociation Studies

LGG was streaked from frozen glycerol stocks onto BHIS agar plates and incubated at Single LGG colonies were used to inoculate 5mL of BHIS broth and grown overnight under static conditions at 37°C in an anaerobic chamber (Coy Laboratory Products, Inc., Grass Lake, MI). The next day, cells were passaged 1/50 into fresh BHIS and grown to an O.D. of 0.4-0.6 nm. Cells were pelleted at 10,000 RPM for 20min at 4°C and resuspended in reduced PBS. 8-12 week old GF C57Bl/6 male mice were fed and maintained on RC or HF in flexible film isolators within the UChicago GRAF for four weeks. After 1 week of diet switch, each mouse received 1.74 ×10^8^ CFUs via gavage in 100μL. Colonization was confirmed in fecal pellets. Serial dilutions of both gavage solution as well as fecal pellets from monoassociated mice after 4 weeks were plated onto BHIS agar and incubated at 37°C anaerobically. Mice were colonized with an average of 2.85×10^9^ ± 1.79×10^9^ CFUs/gram of feces.

### REG3γ Bactericidal Assay

Recombinant REG3γ (rREG3γ) was prepared as previously described (Cash et al., 2006b) and stored at −80°C. *E. faecalis, E. coli K12*, LGG, *Peptostreptococcaceae*, *Bacteroides Thetaiotamicron, L. reuteri, L. intestinalis, L. murinus*, and *L. johnsonii* were re-inoculated into fresh media and incubated at 37°C to mid-log phase. Cultures were spun, resuspended in standard assay buffer (10mM MES pH6, 25mM NaCl), re-pelleted, and diluted 1:25 in standard assay buffer. rREG3γ (0μM, 5μM, 10μM, 20μM) was added followed by incubation at 37°C for 2hrs. After 2hr, dilution plating (1:10 and 1:100) was performed on agar media plates and incubated at 37°C for 24-48hrs under aerobic or anaerobic conditions. Colony forming units (CFUs) were determined and the average % remaining bacteria was calculated relative to 0μM CFUs for each respective bacterial strain.

### Minimum Inhibitory Concentration (MIC) Assay

For the selected *Lactobacillus* strains, *E. faecalis*, *E. coli K12*, *P. stomatis*, *L. monocytogenes*, *B. thetaiotamicron*, and *P. anaerobius*, the minimal inhibitory concentration (MIC) (μM/mL) for rREG3γ was determined. Bacterial cultures in the exponential growth phase were diluted to a turbidity of 0.2 +/− 0.02 and diluted 1:100 in each bacteria-specific media. 50ul of each diluted inoculum was added to each well of 96-well microdilution plate containing 50μl rREG3γ (at concentrations of 15 – 0.01μM) or vehicle control. Inoculated plates were incubated aerobically/anaerobically at 37°C for 24hrs. After incubation, a 96-well plate reader was used to determine O.D. at 600nm. MIC was determined as the lowest concentration of REG3γ that reduced O.D. by 50% or 90% for MIC50 or MIC90.

### Intraperitoneal Glucose Tolerance Test

Male 8-10 week old SPF *Reg3γ*^+/−^, or *Reg3γ*^−/−^ mice on a C57Bl/6 background fed RC or HF were individually housed for 3 weeks. Mice were fasted overnight for 12hrs. A 20% glucose in water solution (2 g/kg body weight) was administered by intraperitoneal (IP) injection. Tail vein blood glucose was measured before and after IP injection at 0, 15, 30, 60, and 120 minutes using a OneTouch Glucose Meter (ADW Diabetes, Pompano Beach, FL). Area under the curve (AUC) was calculated and compared between genotypes using GraphPad Prism v8.

## Quantification and Statistical Analysis

Data from *in vivo* and *in vitro* studies are presented as mean ± SEM or box and whisker plots as mean ± min/max. Statistical analysis was performed using two-way Analysis of Variance (ANOVA) tests followed by Tukey’s test; Brown-Forsythe and Welch ANOVA followed by Dunnett’s test; or Welch’s t test; *p*<0.05 was considered significant. Significant changes in OTU abundances were assessed using ANOVA, as implemented in QIIME (Bonferroni correction for multiple tests; α=0.05). Multivariate statistical tests run on microbial community structure data include Analysis of Similarity (ANOSIM) and adonis tests, as implemented in QIIME. Relative abundances of individual OTUs over the course of 24hrs were compared against the relative expression of host expression of *Reg3γ* in the ileum of both RC and HF SPF mice using Pearson Linear Regression correlation analysis and Spearman RHO non-parametric correlation analysis. False Discovery Rate (FDR) and Bonferroni were used as multi-test correction factors to correct the p-values to reduce Type I statistical error. Significant circadian oscillation of gene expression or copy number was determined using the software CircWave V1.4; a rhythm was determined as present by *p*<0.05. Significant microbial OTU abundance oscillations were determined via e_JTK_Cycle; an OTU was determined rhythmic at GammaP<0.05 (Hutchison et al., 2015).

## Data and Software Availability

The 16S rRNA amplicon raw sequencing files have been deposited in the NCBI Sequence Read Archive under ID code SRP187770.

## KEY RESOURCES TABLE

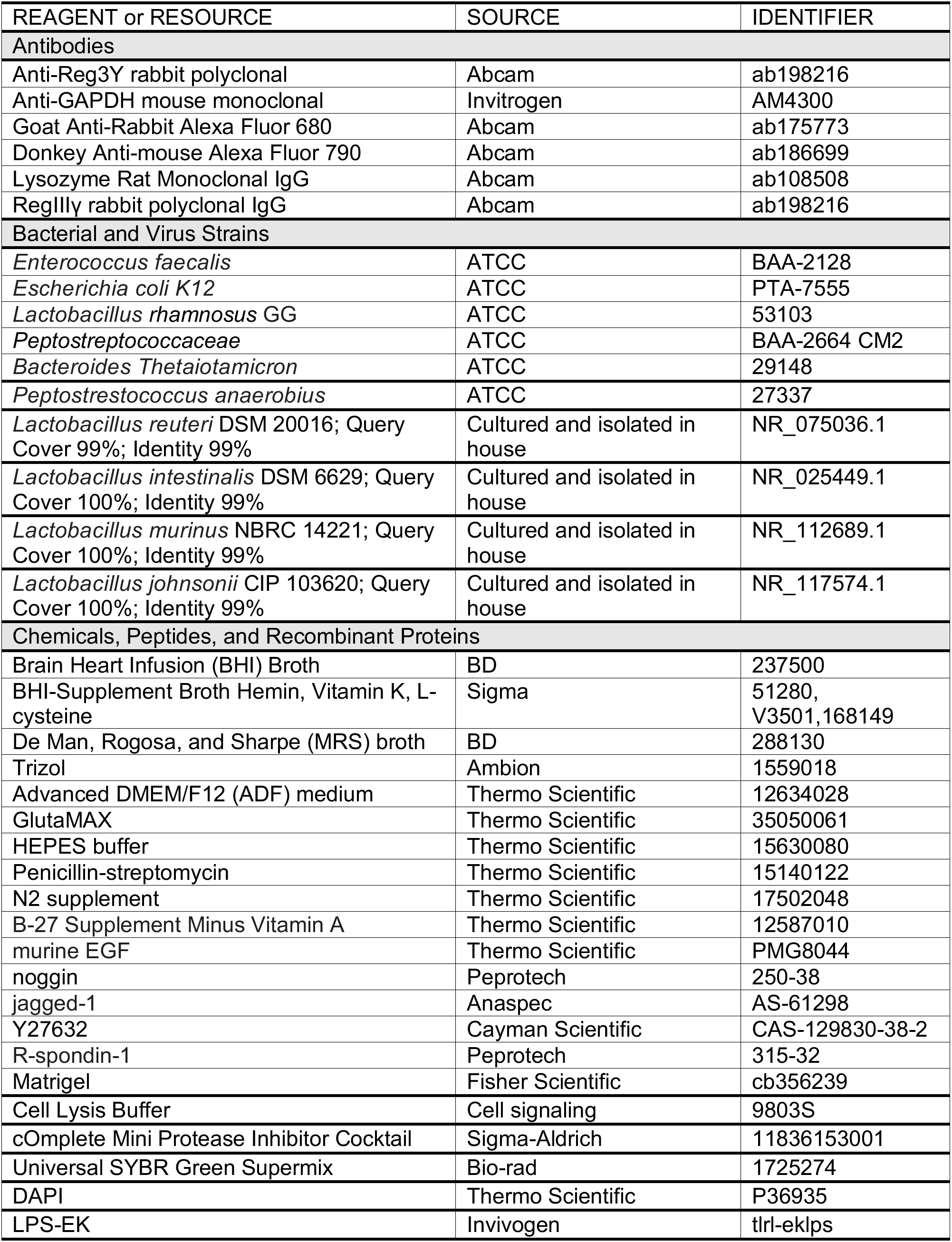
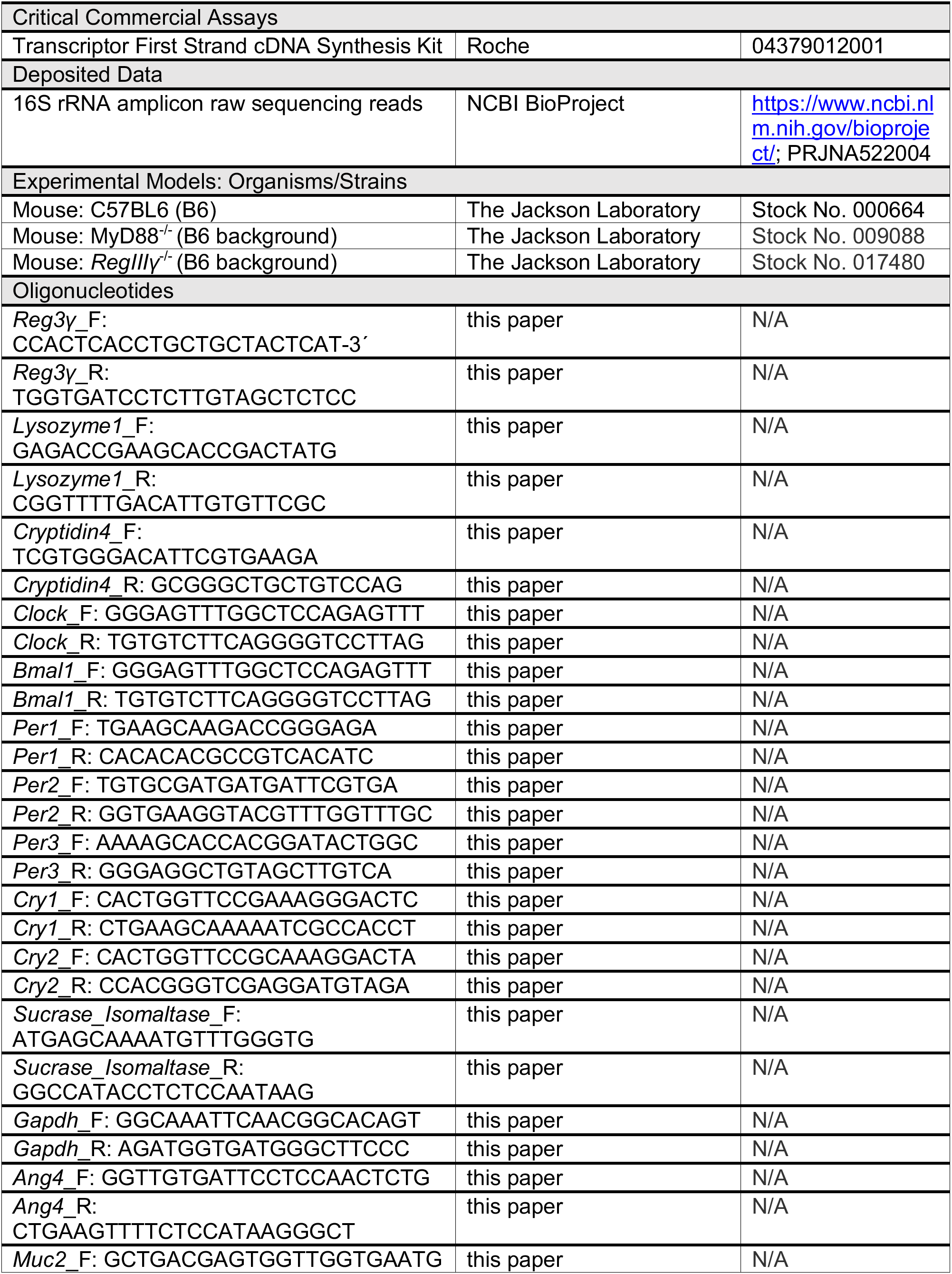
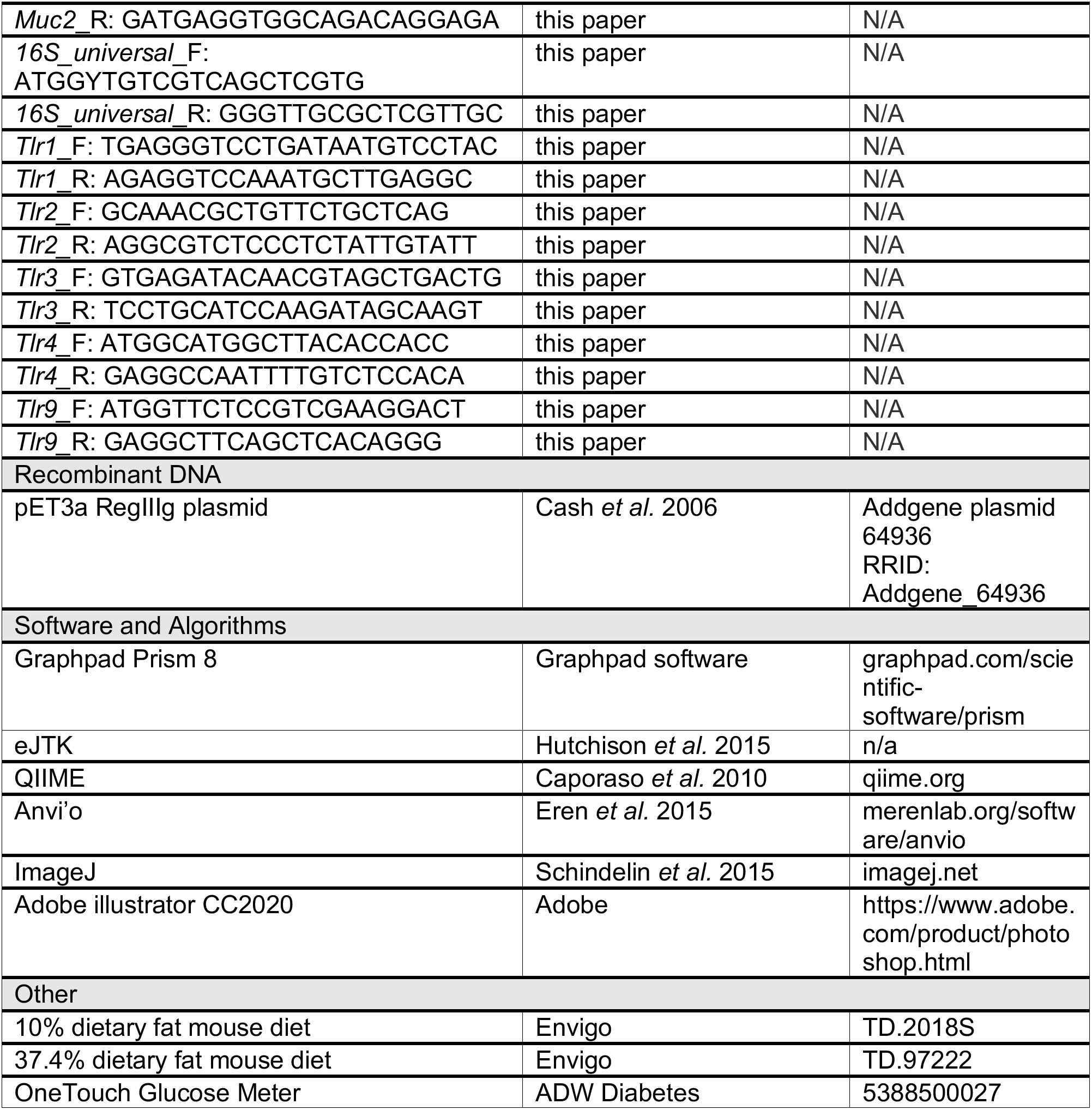

---

**Table S1.** CircWave Co-sinor p-values. **Related to Figures 1, 2, 4, 5, S1, S2, S4**.

|  | Microbial status |  |  |  |
| --- | --- | --- | --- | --- |
|  | SPF | GF | SPF | GF |
|  | Diet |  |  |  |
|  | RC |  | HF |  |
| Gene target | CircWave p-value* |  |  |  |
| *Bmal1* | 6.00E-07 | 0.0001 | 0.0001 | 0.0006 |
| *Clock* | 0.008 | n.s. | 0.007 | n.s. |
| *Per1* | n.s. | 0.009 | 0.025 | n.s. |
| *Per2* | 0.01 | 0.0002 | 0.01 | 0.008 |
| *Per3* | 0.02 | 0.0002 | 0.0005 | 0.0002 |
| *Cry1* | 0 | n.s. | 4.30E-06 | 0.02 |
| *Cry2* | 0.0006 | n.s. | 0.0085 | 0.048 |
| *Reg3y* | 0.04 | n.s. | 0.0009 | n.s. |
| *Lyz1* | n.s. | n.s. | n.s. | n.s. |
| *Ang4* | n.s. | n.s. | n.s. | n.s. |
| *Crypt4* | n.s. | n.s. | n.s. | n.s. |
| *Muc2* | n.s. | 0.003 | n.s. | n.s. |
| 16S rRNA<br>gene copy # | n.s. | ----- | n.s. | ----- |
|  | Diet |  |  |  |
|  | RC | HF | RC | HF |
|  | Genotype |  |  |  |
|  | *Reg3y*<sup>+/-</sup> |  | *Reg3y*<sup>-/-</sup> |  |
| Gene target | CircWave p-value* |  |  |  |
| *Bmal1* | n.s. | n.s. | 0.0013 | 0.0014 |
| *Clock* | n.s. | n.s. | n.s. | 0.0009 |
| *Per1* | n.s. | n.s. | n.s. | 0.032 |
| *Per2* | n.s. | n.s. | n.s. | n.s. |
| *Per3* | n.s. | 0.0159 | n.s. | 0.0037 |
| *Cry1* | 0.0228 | 0.0219 | n.s. | n.s. |
| *Cry2* | n.s. | n.s. | n.s. | n.s. |
| *Reg3y* | 1.70E-06 | n.s. | n.s. | n.s. |
| *Lyz1* | n.s. | n.s. | 0.0469 | n.s. |
| *Ang4* | n.s. | n.s. | n.s. | n.s. |
| *Crypt4* | n.s. | n.s. | n.s. | n.s. |
| *Muc2* | 2.28E-05 | n.s. | n.s. | n.s. |
| 16S rRNA<br>gene copy # | n.s. | n.s. | 0.0367 | n.s. |

\*Significant co-sinor expression pattern indicated by $p < 0.05$
n.s. = Not significant

**Table S2.** Statistical analysis of OTU relative abundances in DILC from RC and HF-fed WT mice determined via ANOVA. OTUs exhibiting Bonferroni P-value less than 0.05 are shown. **Related to Figures 2, S2**.

| Taxonomy | OTU ID | Test-stat | FDR P-value | Bonferroni P-value | RC mean relative abundance | HF mean relative abundance |
| --- | --- | --- | --- | --- | --- | --- |
| p__Firmicutes; c__Clostridia; o__Clostridiales; f__Clostridiaceae | 270200 | 91.2795535 | 1.15E-07 | 1.15E-07 | 0.06666667 | 8.5 |
| p__Firmicutes; c__Clostridia; o__Clostridiales | 270382 | 2.14E-08 | 4.44E-06 | 1.33E-05 | 0.13333333 | 17.125 |
| p__Firmicutes; c__Clostridia; o__Clostridiales; f__Clostridiaceae | 272964 | 42.5045951 | 4.03E-05 | 0.00024152 | 0 | 11.125 |
| p__Firmicutes; c__Clostridia; o__Clostridiales; f__Clostridiaceae | 308802 | 40.5376344 | 5.22E-05 | 0.0003651 | 0 | 19.5 |
| p__Firmicutes; c__Clostridia; o__Clostridiales; f__Clostridiaceae | 297143 | 39.8811545 | 5.25E-05 | 0.00042022 | 0 | 13.5 |
| p__Firmicutes; c__Clostridia; o__Clostridiales; f__Clostridiaceae | 555945 | 38.0056438 | 6.33E-05 | 0.0006329 | 14.6666667 | 10293.0625 |
| p__Firmicutes; c__Clostridia; o__Clostridiales; f__Peptostreptococcaceae | 299474 | 37.4706922 | 6.48E-05 | 0.00071287 | 0 | 78.6875 |
| p__Firmicutes; c__Clostridia; o__Clostridiales; f__Peptostreptococcaceae | 198209 | 36.5559682 | 6.97E-05 | 0.00087576 | 0 | 18.5 |
| p__Firmicutes; c__Clostridia; o__Clostridiales; f__Peptostreptococcaceae | 180516 | 36.403972 | 6.97E-05 | 0.00090649 | 0 | 6.75 |
| p__Firmicutes; c__Clostridia; o__Clostridiales; f__Clostridiaceae | 189407 | 35.3035256 | 8.33E-05 | 0.0011665 | 0 | 16.1875 |
| p__Firmicutes; c__Clostridia; o__Clostridiales; f__Peptostreptococcaceae | 2658058 | 34.261319 | 9.92E-05 | 0.00148741 | 0 | 22.5625 |
| p__Firmicutes; c__Clostridia; o__Clostridiales; f__Peptostreptococcaceae | 262060 | 33.8324487 | 0.00010182 | 0.00164586 | 0.13333333 | 27.8125 |
| p__Firmicutes; c__Clostridia; o__Clostridiales; f__Clostridiaceae | 4396655 | 33.6200207 | 0.00010182 | 0.00173095 | 0 | 8.3125 |
| p__Firmicutes; c__Clostridia; o__Clostridiales; f__Clostridiaceae | 199268 | 32.5943636 | 0.0001165 | 0.00221353 | 0 | 12.9375 |
| p__Firmicutes; c__Clostridia; o__Clostridiales; f__Peptostreptococcaceae | 4380137 | 31.8914956 | 0.00013132 | 0.00262637 | 0 | 7.5 |
| p__Firmicutes; c__Clostridia; o__Clostridiales; f__Clostridiaceae | 191803 | 30.2844404 | 0.00017017 | 0.0039139 | 0 | 18 |
| p__Firmicutes; c__Clostridia; o__Clostridiales; f__Clostridiaceae | 4238179 | 29.2910551 | 0.00018657 | 0.00503728 | 6.2 | 77.8125 |
| p__Firmicutes; c__Clostridia; o__Clostridiales; f__Peptostreptococcaceae | 531374 | 28.4401959 | 0.00020918 | 0.00627536 | 0 | 5.1875 |
| p__Firmicutes; c__Clostridia; o__Clostridiales; f__Peptostreptococcaceae | 514988 | 27.5382615 | 0.0002565 | 0.00795165 | 0.13333333 | 19.625 |
| p__Firmicutes; c__Clostridia; o__Clostridiales | 316496 | 26.8123327 | 0.00030153 | 0.00964901 | 0.13333333 | 22.4375 |
| p__Firmicutes; c__Clostridia; o__Clostridiales; f__Clostridiaceae | 331881 | 26.6387914 | 0.00030636 | 0.01010979 | 0.06666667 | 8.6875 |
| p__Firmicutes; c__Clostridia; o__Clostridiales; f__Peptostreptococcaceae | 322840 | 25.7399593 | 0.00036872 | 0.01290507 | 0 | 28.5625 |
| p__Bacteroidetes; c__Bacteroidia; o__Bacteroidales; f__S24-7 | 211761 | 25.1705115 | 0.00041936 | 0.01509705 | 14.8 | 3.1875 |
| p__Firmicutes; c__Clostridia; o__Clostridiales; f__Peptostreptococcaceae | 1896700 | 24.1234347 | 0.00051895 | 0.02023918 | 0 | 3.875 |
| p__Firmicutes; c__Clostridia; o__Clostridiales | 248126 | 23.2327365 | 0.00060792 | 0.0260987 | 0 | 11.8125 |
| p__Bacteroidetes; c__Bacteroidia; o__Bacteroidales; f__S24-7 | 445144 | 23.0523244 | 0.00060975 | 0.02749365 | 6785.86667 | 1321.375 |
| p__Bacteroidetes; c__Bacteroidia; o__Bacteroidales; f__S24-7 | 331720 | 22.9832477 | 0.00060975 | 0.02804873 | 242.066667 | 64 |
| p__Firmicutes; c__Clostridia; o__Clostridiales; f__Peptostreptococcaceae | 347714 | 22.7885602 | 0.0006251 | 0.02967879 | 0 | 9.9375 |
| p__Firmicutes; c__Clostridia; o__Clostridiales; f__Clostridiaceae; g__Clostridium | 235424 | 21.4259046 | 0.000837 | 0.04436084 | 0.4 | 191.75 |
| p__Firmicutes; c__Clostridia; o__Clostridiales; f__Clostridiaceae | 575768 | 21.3329183 | 0.0008447 | 0.04561388 | 5.06666667 | 18.625 |

**Table S3.** Adonis and ANOSIM R statistics and P-values from Beta-diversity analyses between DILC from RC and HF-fed WT mice, and from DILC and DIMS from RC and HF-fed *Reg3γ*^+/−^ and *Reg3γ*^−/−^ mice. **Related to Figures 2, 5, S2, S5**.

| DILC | Canberra: RC vs HF |  |  |  | Bray Curtis: RC vs HF |  |  |  |
| --- | --- | --- | --- | --- | --- | --- | --- | --- |
|  | adonis |  | ANOSIM |  | adonis |  | ANOSIM |  |
|  | R<sup>2</sup> | P value | R | P value | R<sup>2</sup> | P value | R | P value |
|  | 0.372 | 0.001 | 0.862 | 0.001 | 0.371 | 0.001 | 0.657 | 0.001 |
| DILC | Bray Curtis: RC *Reg3γ<sup>+/-</sup>*, *Reg3γ<sup>-/-</sup>* vs HF *Reg3γ<sup>+/-</sup>*, *Reg3γ<sup>-/-</sup>* |  |  |  |  |  |  |  |
|  | adonis |  |  |  | ANOSIM |  |  |  |
|  | R<sup>2</sup> |  | P value |  | R |  | P value |  |
|  | 0.2253 |  | 0.001 |  | 0.3426 |  | 0.001 |  |
|  | Bray Curtis: RC, *Reg3γ<sup>+/-</sup>* vs *Reg3γ<sup>-/-</sup>* |  |  |  | Bray Curtis: HF, *Reg3γ<sup>+/-</sup>* vs *Reg3γ<sup>-/-</sup>* |  |  |  |
|  | adonis |  | ANOSIM |  | adonis |  | ANOSIM |  |
|  | R<sup>2</sup> | P value | R | P value | R<sup>2</sup> | P value | R | P value |
|  | 0.0653 | 0.003 | 0.1064 | 0.0089 | 0.0234 | 0.328 | 0.0086 | 0.28 |
|  | Bray Curtis: *Reg3γ<sup>+/-</sup>*, RC vs HF |  |  |  | Bray Curtis: *Reg3γ<sup>-/-</sup>*, RC vs HF |  |  |  |
|  | adonis | ANOSIM | adonis | ANOSIM |  |  |  |  |
|  | R<sup>2</sup> | P value | R | P value | R<sup>2</sup> | P value | R | P value |
|  | 0.2127 | 0.001 | 0.532 | 0.001 | 0.194 | 0.001 | 0.4215 | 0.001 |
| DIMS | Bray Curtis: RC *Reg3γ<sup>+/-</sup>*, *Reg3γ<sup>-/-</sup>* vs HF *Reg3γ<sup>+/-</sup>*, *Reg3γ<sup>-/-</sup>* |  |  |  |  |  |  |  |
|  | adonis |  |  |  | ANOSIM |  |  |  |
|  | R<sup>2</sup> |  | P value |  | R |  | P value |  |
|  | 0.1691 |  | 0.001 |  | 0.2786 |  | 0.001 |  |
|  | Bray Curtis: RC, *Reg3γ<sup>+/-</sup>* vs *Reg3γ<sup>-/-</sup>* |  |  |  | Bray Curtis: HF, *Reg3γ<sup>+/-</sup>* vs *Reg3γ<sup>-/-</sup>* |  |  |  |
|  | adonis |  | ANOSIM |  | adonis |  | ANOSIM |  |
|  | R<sup>2</sup> | P value | R | P value | R<sup>2</sup> | P value | R | P value |
|  | 0.032 | 0.136 | 0.0297 | 0.148 | 0.0211 | 0.4 | 0.0015 | 0.38 |
|  | Bray Curtis: *Reg3γ<sup>+/-</sup>*, RC vs HF |  |  |  | Bray Curtis: *Reg3γ<sup>-/-</sup>*, RC vs HF |  |  |  |
|  | adonis |  | ANOSIM |  | adonis |  | ANOSIM |  |
|  | R<sup>2</sup> | P value | R | P value | R<sup>2</sup> | P value | R | P value |
|  | 0.166 | 0.001 | 0.4465 | 0.001 | 0.1408 | 0.001 | 0.3364 | 0.001 |

**Table S4.** eJTK OTU rhythmicity in DILC of RC and HF-fed SPF WT mice. **Related to Figures 2, S2**

| Diet | Taxonomy | OTU_ID | Phase | Mean | Std_Dev | Max | Min | Max_Amp | Tau | P | Gamma_P | Gamma_BH |
| --- | --- | --- | --- | --- | --- | --- | --- | --- | --- | --- | --- | --- |
|  | p < 0.05 |  |  |  |  |  |  |  |  |  |  |  |
| RC | p__Firmicutes; c__Clostridia; o__Clostridiales; f__Clostridiaceae | 225316 | 0 | 0.867 | 1.087 | 3 | 0 | 3 | 0.758 | 4.13E-05 | 0.0006 | 0.081 |
|  | p__Firmicutes; c__Clostridia; o__Clostridiales; f__Lachnospiraceae | 345709 | 16 | 54.067 | 94.339 | 339 | 0 | 339 | 0.736 | 6.52E-05 | 0.0009 | 0.081 |
|  | p__Firmicutes; c__Bacilli; o__Lactobacillales; f__Lactobacillaceae; g__Lactobacillus | 182764 | 0 | 90.4 | 93.203 | 263 | 0 | 263 | 0.718 | 9.45E-05 | 0.0009 | 0.081 |
|  | p__Firmicutes; c__Clostridia; o__Clostridiales | 459276 | 16 | 6.467 | 16.796 | 68 | 0 | 68 | 0.674 | 0.0002 | 0.0035 | 0.081 |
|  | p__Firmicutes; c__Clostridia; o__Clostridiales | 406247 | 20 | 2.067 | 2.977 | 10 | 0 | 10 | 0.672 | 0.0002 | 0.0036 | 0.081 |
|  | p__Firmicutes; c__Clostridia; o__Clostridiales | 1108078 | 16 | 3.4 | 5.401 | 18 | 0 | 18 | 0.637 | 0.0005 | 0.0039 | 0.081 |
|  | p__Firmicutes; c__Clostridia; o__Clostridiales; f__Clostridiaceae | 256232 | 0 | 0.933 | 1.526 | 5 | 0 | 5 | 0.661 | 0.0003 | 0.0039 | 0.081 |
|  | p__Firmicutes; c__Clostridia; o__Clostridiales; f__Lachnospiraceae; g__Coprococcus | 1105328 | 12 | 0.667 | 1.135 | 3 | 0 | 3 | 0.629 | 0.0005 | 0.0039 | 0.081 |
|  | p__Firmicutes; c__Clostridia; o__Clostridiales | 340650 | 16 | 0.8 | 1.833 | 7 | 0 | 7 | 0.637 | 0.0005 | 0.0039 | 0.081 |
|  | p__Firmicutes; c__Clostridia; o__Clostridiales | 346719 | 16 | 10 | 19.610 | 70 | 0 | 70 | 0.637 | 0.0005 | 0.0039 | 0.081 |
|  | p__Firmicutes; c__Clostridia; o__Clostridiales; f__Ruminococcaceae | 375106 | 16 | 0.933 | 1.769 | 7 | 0 | 7 | 0.630 | 0.0005 | 0.0039 | 0.081 |
|  | p__Firmicutes; c__Clostridia; o__Clostridiales | 400599 | 16 | 1.333 | 3.360 | 13 | 0 | 13 | 0.637 | 0.0005 | 0.0039 | 0.081 |
|  | p__Firmicutes; c__Clostridia; o__Clostridiales; f__Lachnospiraceae | 355175 | 16 | 2.933 | 4.509 | 16 | 0 | 16 | 0.654 | 0.0003 | 0.0039 | 0.081 |
|  | p__Firmicutes; c__Clostridia; o__Clostridiales; f__Ruminococcaceae | 346804 | 16 | 0.667 | 1.491 | 6 | 0 | 6 | 0.621 | 0.0006 | 0.0039 | 0.081 |
|  | p__Firmicutes; c__Clostridia; o__Clostridiales; f__Ruminococcaceae; g__Ruminococcus | 278098 | 20 | 2 | 2.658 | 8 | 0 | 8 | 0.634 | 0.0005 | 0.0039 | 0.081 |
|  | p__Firmicutes; c__Clostridia; o__Clostridiales; f__Lachnospiraceae | 355771 | 16 | 9.4 | 22.633 | 81 | 0 | 81 | 0.629 | 0.0005 | 0.0039 | 0.081 |
|  | p__Firmicutes; c__Clostridia; o__Clostridiales | 307170 | 16 | 3.533 | 7.890 | 31 | 0 | 31 | 0.654 | 0.0003 | 0.0039 | 0.081 |
|  | p__Firmicutes; c__Clostridia; o__Clostridiales | 335267 | 16 | 0.933 | 1.948 | 7 | 0 | 7 | 0.631 | 0.0005 | 0.0039 | 0.081 |
|  | p__Firmicutes; c__Clostridia; o__Clostridiales | 1571092 | 16 | 7.867 | 16.095 | 63 | 0 | 63 | 0.611 | 0.0008 | 0.0079 | 0.114 |
|  | p__Firmicutes; c__Clostridia; o__Clostridiales | 826541 | 16 | 2.933 | 3.415 | 10 | 0 | 10 | 0.615 | 0.0007 | 0.0079 | 0.114 |
|  | p__Firmicutes; c__Bacilli; o__Turicibacterales; f__Turicibacteraceae; g__Turicibacter | 347529 | 4 | 2.133 | 1.310 | 4 | 0 | 4 | 0.605 | 0.0008 | 0.0079 | 0.114 |
|  | p__Firmicutes; c__Clostridia; o__Clostridiales; f__Ruminococcaceae; g__Oscillospira | 307608 | 20 | 1.067 | 2.323 | 8 | 0 | 8 | 0.598 | 0.0009 | 0.0079 | 0.114 |
|  | p__Tenericutes; c__Mollicutes; o__Anaeroplasmatales; f__Anaeroplasmataceae; g__Anaeroplasma | 252684 | 16 | 1.467 | 4.485 | 18 | 0 | 18 | 0.598 | 0.0009 | 0.0079 | 0.114 |
|  | p__Firmicutes; c__Clostridia; o__Clostridiales; f__Lachnospiraceae | 344726 | 0 | 6.267 | 9.692 | 33 | 0 | 33 | 0.579 | 0.0013 | 0.0129 | 0.129 |
|  | p__Firmicutes; c__Clostridia; o__Clostridiales | 697874 | 16 | 1.8 | 3.468 | 11 | 0 | 11 | 0.594 | 0.0010 | 0.0129 | 0.129 |
|  | p__Firmicutes; c__Clostridia; o__Clostridiales; f__Lachnospiraceae; g__Anaerostipes | 534926 | 16 | 7.4 | 13.908 | 56 | 0 | 56 | 0.588 | 0.0011 | 0.0129 | 0.129 |
|  | p__Firmicutes; c__Clostridia; o__Clostridiales | 229452 | 16 | 18.667 | 49.076 | 198 | 0 | 198 | 0.579 | 0.0013 | 0.0129 | 0.129 |
|  | p__Firmicutes; c__Clostridia; o__Clostridiales | 1106614 | 16 | 16.667 | 29.236 | 98 | 0 | 98 | 0.585 | 0.0012 | 0.0129 | 0.129 |
|  | p__Firmicutes; c__Clostridia; o__Clostridiales; f__Lachnospiraceae | 305012 | 16 | 3.867 | 7.745 | 30 | 0 | 30 | 0.595 | 0.0010 | 0.0129 | 0.129 |
|  | p__Firmicutes; c__Clostridia; o__Clostridiales | 408877 | 16 | 5.133 | 11.871 | 45 | 0 | 45 | 0.579 | 0.0013 | 0.0129 | 0.129 |
|  | p__Firmicutes; c__Clostridia; o__Clostridiales | 345862 | 16 | 8.333 | 12.451 | 37 | 0 | 37 | 0.578 | 0.0013 | 0.0129 | 0.129 |
|  | p__Firmicutes; c__Clostridia; o__Clostridiales | 356590 | 16 | 15.933 | 21.947 | 75 | 0 | 75 | 0.581 | 0.0013 | 0.0129 | 0.129 |
|  | k__Bacteria; p__Cyanobacteria; c__Chloroplast; o__Streptophyta | 3541 | 0 | 7.133 | 12.516 | 41 | 0 | 41 | 0.561 | 0.0018 | 0.0219 | 0.200 |
|  | p__Firmicutes; c__Clostridia; o__Clostridiales; f__Ruminococcaceae; g__Oscillospira | 328905 | 16 | 0.867 | 1.628 | 6 | 0 | 6 | 0.567 | 0.0016 | 0.0219 | 0.200 |
|  | p__Firmicutes; c__Clostridia; o__Clostridiales; f__Ruminococcaceae | 331737 | 16 | 5.8 | 11.594 | 45 | 0 | 45 | 0.555 | 0.0020 | 0.0292 | 0.259 |
|  | p__Firmicutes; c__Clostridia; o__Clostridiales | 1112121 | 20 | 16.333 | 57.426 | 231 | 0 | 231 | 0.553 | 0.0020 | 0.0302 | 0.260 |
|  | p__Firmicutes; c__Clostridia; o__Clostridiales; f__Lachnospiraceae | 316946 | 16 | 1.4 | 3.460 | 13 | 0 | 13 | 0.548 | 0.0022 | 0.0331 | 0.260 |
|  | p__Firmicutes; c__Clostridia; o__Clostridiales; f__Ruminococcaceae | 170926 | 20 | 1.067 | 2.620 | 10 | 0 | 10 | 0.548 | 0.0022 | 0.0331 | 0.260 |
|  | p__Firmicutes; c__Clostridia; o__Clostridiales; f__Ruminococcaceae | 434089 | 0 | 4.4 | 8.732 | 33 | 0 | 33 | 0.541 | 0.0025 | 0.0359 | 0.260 |
|  | p__Firmicutes; c__Clostridia; o__Clostridiales; f__Ruminococcaceae; g__Oscillospira | 277588 | 16 | 0.467 | 1.258 | 5 | 0 | 5 | 0.538 | 0.0026 | 0.0359 | 0.260 |
|  | p__Firmicutes; c__Clostridia; o__Clostridiales; f__Ruminococcaceae | 1110008 | 16 | 3.533 | 7.374 | 28 | 0 | 28 | 0.541 | 0.0025 | 0.0360 | 0.260 |
|  | p__Firmicutes; c__Clostridia; o__Clostridiales; f__Ruminococcaceae | 318370 | 16 | 2.000 | 4.131 | 16 | 0 | 16 | 0.540 | 0.0025 | 0.0360 | 0.260 |
|  | p__Firmicutes; c__Clostridia; o__Clostridiales | 343581 | 16 | 12.467 | 27.131 | 99 | 0 | 99 | 0.535 | 0.0027 | 0.0408 | 0.269 |
|  | p__Firmicutes; c__Clostridia; o__Clostridiales; f__Lachnospiraceae | 354163 | 20 | 12.800 | 20.582 | 69 | 0 | 69 | 0.534 | 0.0028 | 0.0413 | 0.269 |
|  | p__Firmicutes; c__Clostridia; o__Clostridiales; f__Ruminococcaceae; g__Oscillospira | 345582 | 16 | 1.533 | 3.775 | 14 | 0 | 14 | 0.531 | 0.0029 | 0.0432 | 0.269 |
|  | p__Firmicutes; c__Clostridia; o__Clostridiales; f__Clostridiaceae | 555945 | 12 | 14.667 | 26.810 | 113 | 1 | 113 | 0.531 | 0.0029 | 0.0433 | 0.269 |
|  | p__Firmicutes; c__Clostridia; o__Clostridiales | 382220 | 16 | 7.333 | 20.758 | 83 | 0 | 83 | 0.529 | 0.0030 | 0.0447 | 0.269 |
|  | p__Firmicutes; c__Clostridia; o__Clostridiales | 377946 | 16 | 1.667 | 3.360 | 11 | 0 | 11 | 0.528 | 0.0030 | 0.0456 | 0.269 |
|  | p__Firmicutes; c__Clostridia; o__Clostridiales; f__Lachnospiraceae; g__Coprococcus | 653533 | 16 | 15.267 | 18.234 | 61 | 0 | 61 | 0.527 | 0.0031 | 0.0464 | 0.269 |
|  | p__Firmicutes; c__Clostridia; o__Clostridiales; f__Ruminococcaceae; g__Oscillospira | 167509 | 16 | 2.667 | 9.714 | 39 | 0 | 39 | 0.519 | 0.0035 | 0.0499 | 0.269 |
|  | p_Firmicutes; c_Clostridia; o_Clostridiales | 747987 | 16 | 1.133 | 2.276 | 8 | 0 | 8 | 0.518 | 0.0035 | 0.0499 | 0.269 |
|  | p_Firmicutes; c_Clostridia; o_Clostridiales | 197290 | 16 | 1.200 | 2.663 | 10 | 0 | 10 | 0.522 | 0.0033 | 0.0499 | 0.269 |
|  | p_Firmicutes; c_Clostridia; o_Clostridiales; f_Ruminococcaceae; g_Oscillospira | 327808 | 16 | 0.733 | 1.123 | 3 | 0 | 3 | 0.517 | 0.0036 | 0.0499 | 0.269 |
| HF | p_Firmicutes; c_Erysipelotrichi; o_Erysipelotrichales; f_Erysipelotrichaceae | 167420 | 0 | 3.875 | 7.432 | 28 | 0 | 28 | 0.684 | 0.0001 | 0.0016 | 0.444 |
|  | p_Firmicutes; c_Bacilli; o_Lactobacillales; f_Lactobacillaceae; g_Lactobacillus | 306306 | 12 | 2.938 | 3.030 | 11 | 0 | 11 | 0.582 | 0.0008 | 0.0124 | 0.693 |
|  | p_Bacteroidetes; c_Bacteroidia; o_Bacteroidales; f_S24-7 | 271131 | 0 | 1.875 | 2.058 | 6 | 0 | 6 | 0.552 | 0.0014 | 0.0214 | 0.693 |
|  | p_Firmicutes; c_Clostridia; o_Clostridiales; f_Ruminococcaceae; g_Oscillospira | 277588 | 16 | 1.438 | 4.808 | 20 | 0 | 20 | 0.530 | 0.0021 | 0.0312 | 0.693 |
|  | p_Firmicutes; c_Clostridia; o_Clostridiales; f_Lachnospiraceae | 352559 | 0 | 5.125 | 12.639 | 53 | 0 | 53 | 0.508 | 0.0030 | 0.0453 | 0.693 |
|  | p_Firmicutes; c_Clostridia; o_Clostridiales; f_Lachnospiraceae | 345536 | 4 | 1.000 | 2.208 | 9 | 0 | 9 | 0.503 | 0.0033 | 0.0495 | 0.693 |
| p = 0.05 – 0.1 |  |  |  |  |  |  |  |  |  |  |  |  |
| RC | p_Bacteroidetes; c_Bacteroidia; o_Bacteroidales; f_S24-7 | 445144 | 16 | 6785.867 | 4072.125 | 12745 | 103 | 12642 | 0.516 | 0.0037 | 0.0549 | 0.285 |
|  | p_Firmicutes; c_Erysipelotrichi; o_Erysipelotrichales; f_Erysipelotrichaceae | 167420 | 16 | 6.733 | 6.486 | 21 | 0 | 21 | 0.516 | 0.0037 | 0.0551 | 0.285 |
|  | p_Firmicutes; c_Clostridia; o_Clostridiales | 348404 | 20 | 4.933 | 7.602 | 28 | 0 | 28 | 0.516 | 0.0037 | 0.0554 | 0.285 |
|  | p_Firmicutes; c_Clostridia; o_Clostridiales | 628218 | 16 | 2.200 | 7.204 | 29 | 0 | 29 | 0.515 | 0.0037 | 0.0561 | 0.285 |
|  | p_Firmicutes; c_Clostridia; o_Clostridiales; f_Ruminococcaceae; g_Oscillospira | 276985 | 16 | 4.667 | 10.593 | 42 | 0 | 42 | 0.510 | 0.0040 | 0.0594 | 0.290 |
|  | p_Firmicutes; c_Clostridia; o_Clostridiales; f_Ruminococcaceae; g_Oscillospira | 1110378 | 0 | 7.667 | 15.023 | 59 | 0 | 59 | 0.503 | 0.0045 | 0.0645 | 0.290 |
|  | p_Firmicutes; c_Clostridia; o_Clostridiales; f_Clostridiaceae | 3358082 | 0 | 1.000 | 1.317 | 4 | 0 | 4 | 0.498 | 0.0049 | 0.0659 | 0.290 |
|  | p_Firmicutes; c_Bacilli; o_Lactobacillales; f_Lactobacillaceae; g_Lactobacillus | 549756 | 20 | 1.200 | 1.869 | 6 | 0 | 6 | 0.498 | 0.0048 | 0.0659 | 0.290 |
|  | p_Firmicutes; c_Bacilli; o_Bacillales; f_Staphylococcaceae; g_Staphylococcus; s_sciuri | 1084865 | 12 | 6.467 | 9.763 | 33 | 0 | 33 | 0.500 | 0.0047 | 0.0659 | 0.290 |
|  | p_Firmicutes; c_Clostridia; o_Clostridiales; f_Lachnospiraceae | 262104 | 16 | 1.533 | 3.442 | 12 | 0 | 12 | 0.499 | 0.0048 | 0.0659 | 0.290 |
|  | p_Firmicutes; c_Clostridia; o_Clostridiales | 194662 | 16 | 3.733 | 10.408 | 42 | 0 | 42 | 0.499 | 0.0048 | 0.0659 | 0.290 |
|  | p_Firmicutes; c_Clostridia; o_Clostridiales; f_Lachnospiraceae | 258202 | 12 | 2.467 | 3.538 | 11 | 0 | 11 | 0.495 | 0.0050 | 0.0702 | 0.302 |
|  | p_Firmicutes; c_Clostridia; o_Clostridiales | 309054 | 16 | 2.067 | 3.820 | 14 | 0 | 14 | 0.495 | 0.0050 | 0.0703 | 0.302 |
|  | p_Firmicutes; c_Bacilli; o_Lactobacillales; f_Lactobacillaceae; g_Lactobacillus | 590168 | 12 | 3.067 | 4.864 | 18 | 0 | 18 | 0.492 | 0.0053 | 0.0730 | 0.308 |
|  | p_Firmicutes; c_Clostridia; o_Clostridiales | 327284 | 16 | 4.933 | 12.678 | 51 | 0 | 51 | 0.490 | 0.0054 | 0.0743 | 0.309 |
|  | p_Firmicutes; c_Clostridia; o_Clostridiales | 564961 | 16 | 0.867 | 2.729 | 11 | 0 | 11 | 0.488 | 0.0056 | 0.0764 | 0.309 |
|  | p_Firmicutes; c_Clostridia; o_Clostridiales; f_Ruminococcaceae | 343140 | 16 | 8.933 | 16.723 | 66 | 0 | 66 | 0.488 | 0.0056 | 0.0766 | 0.309 |
|  | p_Firmicutes; c_Clostridia; o_Clostridiales; f_Lachnospiraceae | 759751 | 16 | 11.067 | 23.245 | 92 | 0 | 92 | 0.484 | 0.0060 | 0.0801 | 0.309 |
|  | p_Firmicutes; c_Clostridia; o_Clostridiales; f_Lachnospiraceae | 1076587 | 20 | 4.933 | 7.818 | 31 | 0 | 31 | 0.484 | 0.0060 | 0.0801 | 0.309 |
|  | p_Firmicutes; c_Clostridia; o_Clostridiales; f_Ruminococcaceae | 275218 | 16 | 1.933 | 5.285 | 21 | 0 | 21 | 0.482 | 0.0062 | 0.0820 | 0.309 |
|  | p_Proteobacteria; c_Alphaproteobacteria; o_Rickettsiales; f_mitochondria | 1892252 | 16 | 9.533 | 24.857 | 100 | 0 | 100 | 0.481 | 0.0062 | 0.0825 | 0.309 |
|  | p_Firmicutes; c_Clostridia; o_Clostridiales | 434339 | 16 | 2.533 | 4.064 | 16 | 0 | 16 | 0.481 | 0.0062 | 0.0826 | 0.309 |
|  | p_Firmicutes; c_Clostridia; o_Clostridiales; f_Ruminococcaceae; g_Oscillospira | 180235 | 16 | 1.200 | 1.869 | 6 | 0 | 6 | 0.480 | 0.0063 | 0.0834 | 0.310 |
|  | p_Firmicutes; c_Clostridia; o_Clostridiales | 339718 | 20 | 2.400 | 6.888 | 28 | 0 | 28 | 0.476 | 0.0067 | 0.0869 | 0.313 |
|  | p_Firmicutes; c_Clostridia; o_Clostridiales | 351062 | 20 | 7.000 | 11.736 | 44 | 0 | 44 | 0.476 | 0.0067 | 0.0869 | 0.313 |
|  | p_Firmicutes; c_Clostridia; o_Clostridiales | 336276 | 16 | 7.200 | 12.357 | 43 | 0 | 43 | 0.475 | 0.0068 | 0.0879 | 0.313 |
|  | p_Firmicutes; c_Clostridia; o_Clostridiales | 351859 | 20 | 6.000 | 8.989 | 26 | 0 | 26 | 0.475 | 0.0068 | 0.0879 | 0.313 |
|  | p_Firmicutes; c_Clostridia; o_Clostridiales; f_Clostridiaceae | 342504 | 16 | 0.467 | 0.618 | 2 | 0 | 2 | 0.472 | 0.0070 | 0.0905 | 0.315 |
|  | p_Firmicutes; c_Clostridia; o_Clostridiales; f_Ruminococcaceae; g_Ruminococcus | 327900 | 20 | 4.267 | 9.896 | 38 | 0 | 38 | 0.472 | 0.0070 | 0.0905 | 0.315 |
|  | p_Firmicutes; c_Clostridia; o_Clostridiales | 272949 | 16 | 9.867 | 28.234 | 113 | 0 | 113 | 0.469 | 0.0074 | 0.0937 | 0.321 |
|  | p_Firmicutes; c_Clostridia; o_Clostridiales | 458550 | 0 | 11.133 | 10.831 | 36 | 0 | 36 | 0.467 | 0.0077 | 0.0964 | 0.326 |
| HF | p_Firmicutes; c_Clostridia; o_Clostridiales; f_Ruminococcaceae; g_Oscillospira | 408513 | 16 | 22.125 | 39.141 | 159 | 1 | 158 | 0.496 | 0.0037 | 0.0556 | 0.693 |
|  | p_Firmicutes; c_Clostridia; o_Clostridiales | 356590 | 16 | 0.375 | 0.696 | 2 | 0 | 2 | 0.485 | 0.0044 | 0.0660 | 0.693 |
|  | p_Firmicutes; c_Clostridia; o_Clostridiales; f_Lachnospiraceae | 355175 | 16 | 0.125 | 0.331 | 1 | 0 | 1 | 0.477 | 0.0050 | 0.0744 | 0.693 |
|  | p_Firmicutes; c_Clostridia; o_Clostridiales | 461487 | 16 | 0.188 | 0.527 | 2 | 0 | 2 | 0.469 | 0.0056 | 0.0847 | 0.693 |
|  | p_Firmicutes; c_Clostridia; o_Clostridiales; f_Lachnospiraceae | 355822 | 16 | 0.313 | 0.982 | 4 | 0 | 4 | 0.469 | 0.0056 | 0.0847 | 0.693 |
|  | p_Firmicutes; c_Clostridia; o_Clostridiales | 346764 | 20 | 0.750 | 2.437 | 10 | 0 | 10 | 0.469 | 0.0056 | 0.0847 | 0.693 |
|  | p_Firmicutes; c_Clostridia; o_Clostridiales; f_Ruminococcaceae; g_Ruminococcus | 278098 | 20 | 0.563 | 0.788 | 2 | 0 | 2 | 0.468 | 0.0058 | 0.0863 | 0.693 |
|  | p_Firmicutes; c_Clostridia; o_Clostridiales | 458550 | 12 | 0.625 | 1.364 | 5 | 0 | 5 | 0.467 | 0.0058 | 0.0873 | 0.693 |
|  | p_Firmicutes; c_Clostridia; o_Clostridiales; f_Lachnospiraceae | 457614 | 16 | 1.375 | 1.798 | 7 | 0 | 7 | 0.466 | 0.0059 | 0.0889 | 0.693 |
|  | p_Firmicutes; c_Clostridia; o_Clostridiales; f_Lachnospiraceae; g_Dorea | 724472 | 16 | 0.813 | 1.424 | 4 | 0 | 4 | 0.465 | 0.0060 | 0.0900 | 0.693 |

**Table S5.** Statistical analysis of OTU relative abundances in DILC from *Reg3γ*^+/−^ vs. *Reg3γ*^−/−^ mice fed RC or HF diet determined via ANOVA. OTUs exhibiting Bonferroni P-values less than 0.05 are shown. **Related to Figures 5, S5**.

| Taxonomy | OTU ID | Test-stat | FDR P-value | Bonferroni P-value | *Reg3γ<sup>+/-</sup>* RC mean relative abundance | *Reg3γ<sup>+/-</sup>* HF mean relative abundance | *Reg3γ<sup>-/-</sup>* RC mean relative abundance | *Reg3γ<sup>-/-</sup>* HF mean relative abundance |
| --- | --- | --- | --- | --- | --- | --- | --- | --- |
| p_Firmicutes; c_Clostridia; o_Clostridiales; f_Clostridiaceae | 555945 | 20.247 | 4.39E-07 | 4.39E-07 | 47.04 | 660.68 | 8.67 | 385.55 |
| p_Firmicutes; c_Bacilli; o_Lactobacillales; f_Streptococcaceae; g_Lactococcus | 716006 | 16.191 | 8.23E-06 | 1.82E-05 | 50.04 | 1117.48 | 14.08 | 933.09 |
| p_Firmicutes; c_Clostridia; o_Clostridiales; f_Peptostreptococcaceae | 198209 | 15.642 | 8.23E-06 | 3.08E-05 | 0.17 | 5.84 | 0.17 | 3.82 |
| p_Firmicutes; c_Clostridia; o_Clostridiales; f_Peptostreptococcaceae | 180516 | 15.497 | 8.23E-06 | 3.54E-05 | 0.48 | 6.6 | 0.17 | 3.14 |
| p_Firmicutes; c_Clostridia; o_Clostridiales; f_Clostridiaceae | 191803 | 15.342 | 8.23E-06 | 4.12E-05 | 0.13 | 1.8 | 0 | 0.45 |
| p_Firmicutes; c_Bacilli; o_Lactobacillales; f_Streptococcaceae; g_Lactococcus | 4468805 | 14.900 | 1.06E-05 | 6.34E-05 | 0.26 | 4.92 | 0 | 4.73 |
| p_Firmicutes; c_Clostridia; o_Clostridiales | 248126 | 14.315 | 1.61E-05 | 0.00011 | 0.04 | 2 | 0 | 1 |
| p_Firmicutes; c_Bacilli; o_Lactobacillales; f_Streptococcaceae; g_Lactococcus | 513080 | 14.065 | 1.64E-05 | 0.00014 | 0.26 | 6.96 | 0.17 | 5.55 |
| p_Proteobacteria; c_Betaproteobacteria; o_Burkholderiales; f_Alcaligenaceae; g_Sutterella | 359809 | 14.047 | 1.64E-05 | 0.00015 | 12.52 | 0.68 | 32.79 | 3.95 |
| p_Firmicutes; c_Clostridia; o_Clostridiales; f_Clostridiaceae | 199268 | 13.512 | 2.52E-05 | 0.00025 | 0.04 | 1.56 | 0.04 | 0.86 |
| p_Firmicutes; c_Clostridia; o_Clostridiales; f_Peptostreptococcaceae | 347714 | 13.172 | 3.10E-05 | 0.00036 | 0.13 | 1.68 | 0 | 1.09 |
| p_Firmicutes; c_Bacilli; o_Lactobacillales; f_Streptococcaceae; g_Lactococcus | 571744 | 13.068 | 3.10E-05 | 0.00040 | 0.13 | 2.08 | 0 | 2.27 |
| p_Firmicutes; c_Bacilli; o_Lactobacillales; f_Streptococcaceae; g_Lactococcus | 1523543 | 13.050 | 3.10E-05 | 0.00040 | 0 | 2.48 | 0 | 2.41 |
| p_Firmicutes; c_Clostridia; o_Clostridiales; f_Peptostreptococcaceae | 514988 | 12.659 | 4.02E-05 | 0.00060 | 0.09 | 2.4 | 0 | 1.41 |
| p_Bacteroidetes; c_Bacteroidia; o_Bacteroidales; f_S24-7 | 270984 | 12.658 | 4.02E-05 | 0.00060 | 14.74 | 1.28 | 27.13 | 4.09 |
| p_Bacteroidetes; c_Bacteroidia; o_Bacteroidales; f_S24-7 | 416078 | 12.443 | 4.71E-05 | 0.00075 | 36.30 | 1 | 41.17 | 7.36 |
| p_Bacteroidetes; c_Bacteroidia; o_Bacteroidales; f_S24-7 | 465480 | 12.118 | 6.07E-05 | 0.00105 | 0.57 | 0 | 2.04 | 0.09 |
| p_Firmicutes; c_Clostridia; o_Clostridiales; f_Peptostreptococcaceae | 178364 | 12.053 | 6.07E-05 | 0.00112 | 0.17 | 1.52 | 0 | 0.64 |
| p_Firmicutes; c_Bacilli; o_Lactobacillales; f_Streptococcaceae; g_Lactococcus | 586387 | 12.033 | 6.07E-05 | 0.00115 | 0.17 | 3.08 | 0 | 2.55 |
| p_Bacteroidetes; c_Bacteroidia; o_Bacteroidales; f_S24-7 | 355746 | 11.715 | 8.03E-05 | 0.00161 | 45.87 | 2.52 | 61.96 | 8.18 |
| p_Firmicutes; c_Clostridia; o_Clostridiales; f_Peptostreptococcaceae | 322840 | 11.517 | 9.42E-05 | 0.00198 | 0.26 | 2.68 | 0 | 1.5 |
| p_Bacteroidetes; c_Bacteroidia; o_Bacteroidales; f_S24-7 | 315430 | 11.258 | 0.0001183 | 0.00260 | 18.61 | 1.04 | 30.21 | 6.23 |
| p_Firmicutes; c_Bacilli; o_Lactobacillales; f_Streptococcaceae; g_Lactococcus | 345575 | 11.116 | 0.0001316 | 0.00303 | 0.17 | 2.44 | 0 | 2.86 |
| p_Firmicutes; c_Clostridia; o_Clostridiales; f_Clostridiaceae | 272964 | 10.653 | 0.0002068 | 0.00496 | 0.13 | 1.16 | 0 | 1.05 |
| p_Firmicutes; c_Clostridia; o_Clostridiales; f_Peptostreptococcaceae | 2658058 | 10.583 | 0.0002143 | 0.00536 | 0.04 | 1.44 | 0.08 | 0.73 |
| p__Bacteroidetes; c__Bacteroidia;<br>o__Bacteroidales; f__S24-7 | 421792 | 10.427 | 0.0002438 | 0.00634 | 89.91 | 2.2 | 106.79 | 16.41 |
| p__Firmicutes; c__Bacilli;<br>o__Lactobacillales;<br>f__Streptococcaceae; g__Lactococcus | 316321 | 10.330 | 0.0002606 | 0.00704 | 0.13 | 2.84 | 0.04 | 2.32 |
| p__Firmicutes; c__Bacilli;<br>o__Lactobacillales;<br>f__Lactobacillaceae; g__Lactobacillus | 604966 | 10.285 | 0.0002637 | 0.00739 | 10.91 | 0.56 | 10.96 | 2 |
| p__Firmicutes; c__Bacilli;<br>o__Lactobacillales;<br>f__Lactobacillaceae; g__Lactobacillus;<br>s__reuteri | 692154 | 10.208 | 0.0002678 | 0.00803 | 15.13 | 0.52 | 18 | 2 |
| p__Bacteroidetes; c__Bacteroidia;<br>o__Bacteroidales; f__S24-7 | 453896 | 10.208 | 0.0002678 | 0.00804 | 32.91 | 2.96 | 57.42 | 9.09 |
| p__Firmicutes; c__Bacilli;<br>o__Lactobacillales;<br>f__Streptococcaceae; g__Lactococcus | 288680 | 9.598 | 0.0005047 | 0.015647 | 0 | 2.2 | 0 | 1.86 |
| p__Firmicutes; c__Clostridia;<br>o__Clostridiales;<br>f__Peptostreptococcaceae | 4380137 | 9.291 | 0.0006861 | 0.02196 | 0 | 0.8 | 0 | 0.23 |
| p__Firmicutes; c__Bacilli;<br>o__Lactobacillales;<br>f__Lactobacillaceae; g__Lactobacillus;<br>s__reuteri | 588197 | 9.229 | 0.0007129 | 0.02353 | 4.04 | 0.52 | 5.54 | 0.86 |
| p__Bacteroidetes; c__Bacteroidia;<br>o__Bacteroidales; f__S24-7 | 195919 | 9.164 | 0.0007435 | 0.02528 | 4.39 | 2.92 | 19 | 6.14 |
| p__Firmicutes; c__Bacilli;<br>o__Lactobacillales;<br>f__Lactobacillaceae; g__Lactobacillus | 1107027 | 8.845 | 0.0010308 | 0.03608 | 530.17 | 134.76 | 101.54 | 161.55 |
| p__Firmicutes; c__Clostridia;<br>o__Clostridiales; f__Clostridiaceae | 189407 | 8.606 | 0.0012965 | 0.04715 | 0.22 | 1.4 | 0 | 0.82 |
| p__Bacteroidetes; c__Bacteroidia;<br>o__Bacteroidales; f__S24-7 | 331720 | 8.591 | 0.0012965 | 0.04797 | 0.74 | 0 | 1.38 | 0.23 |

**Table S6.**
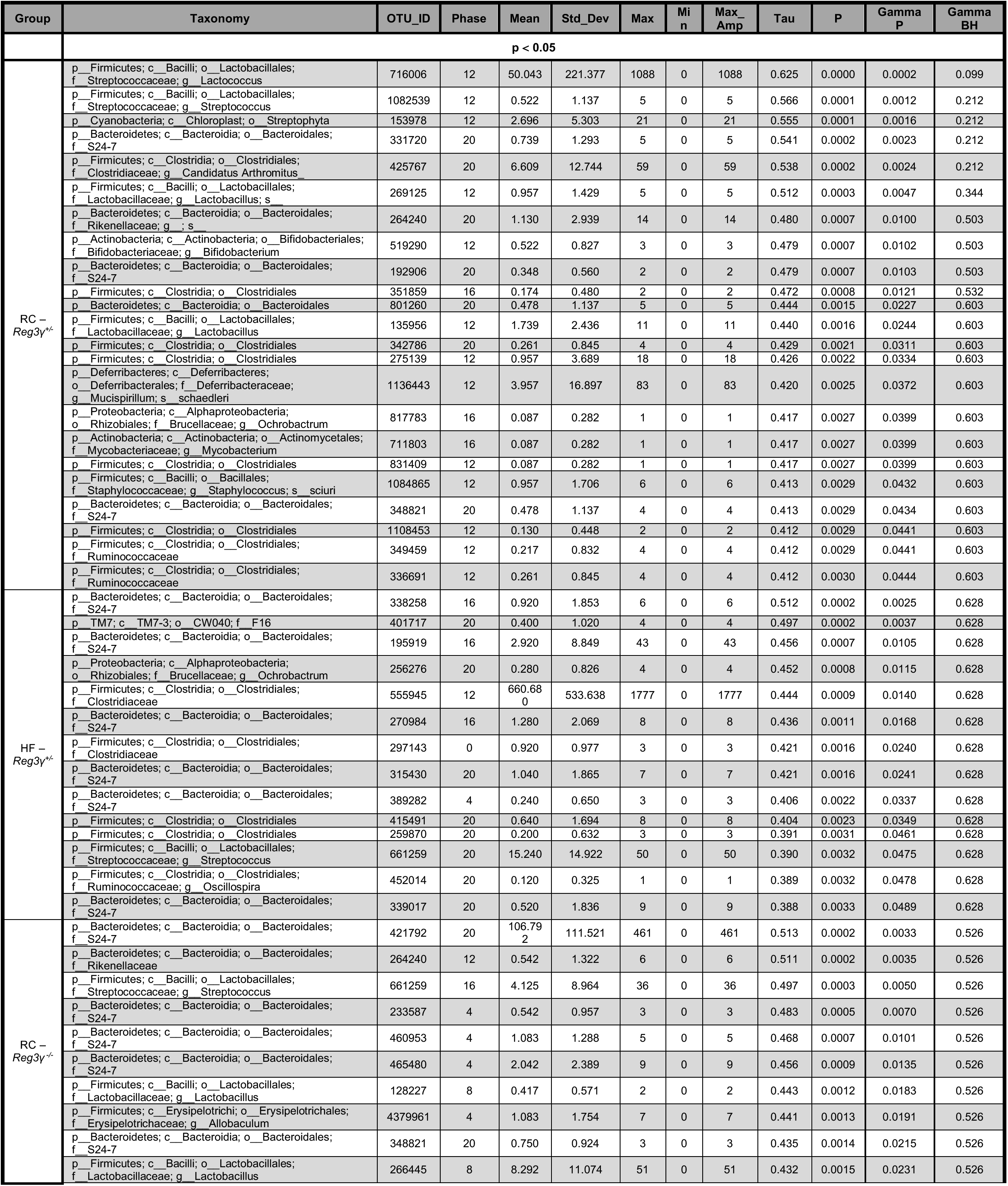
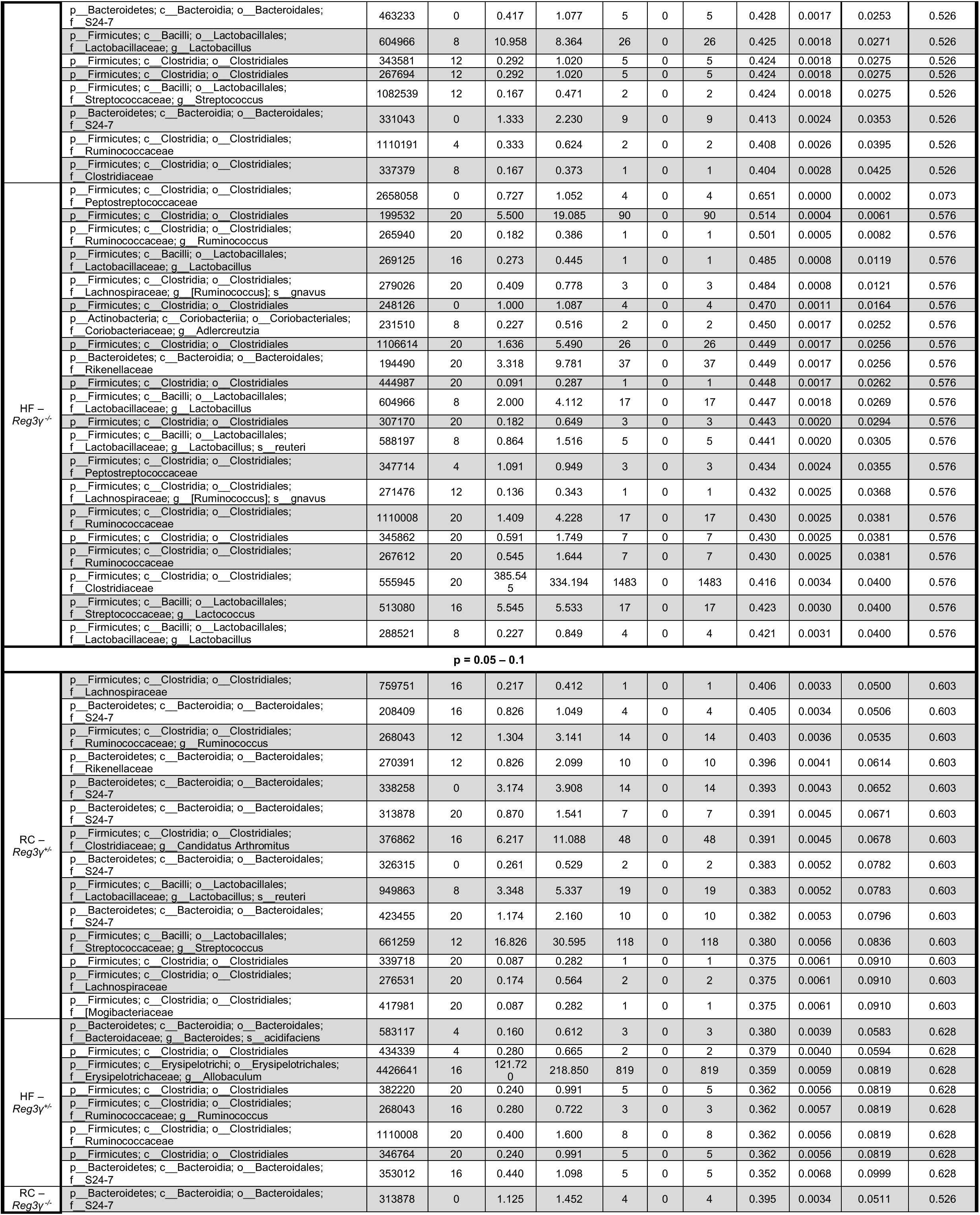
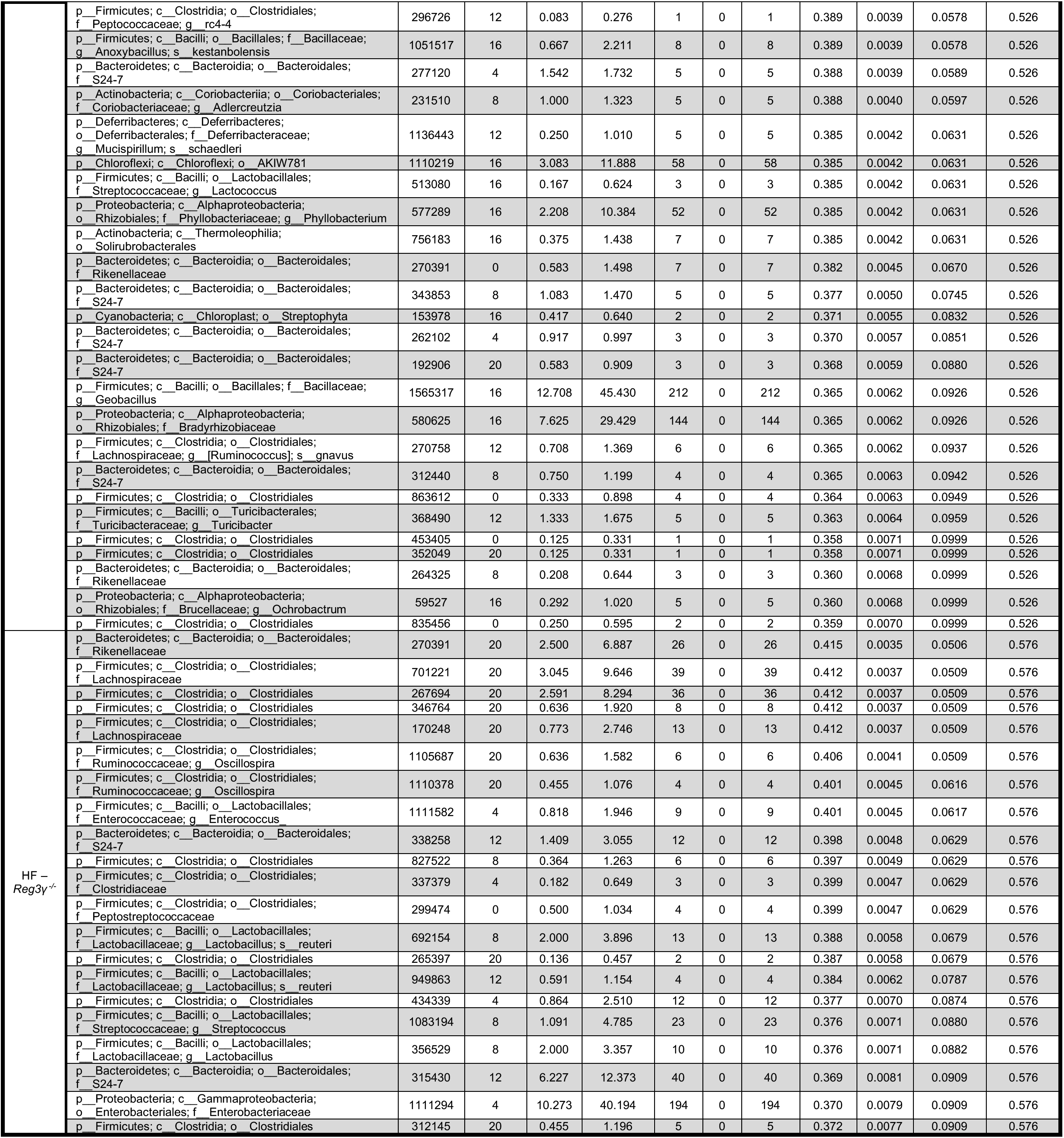
eJTK OTU rhythmicity in DILC from RC and HF-fed *Reg3γ*^+/−^ and *Reg3γ*^−/−^ mice. Related to **Figures 5, 6, S5**

**Table S7.** eJTK oscillating OTUs in DILC and DIMS at the taxonomic levels of order and family in RC and HF-fed *Reg3γ*^+/−^ and *Reg3γ*^−/−^ mice. **Related to Figure 6**.

| Order | Family | DILC |  |  |  | DIMS |  |  |  |
| --- | --- | --- | --- | --- | --- | --- | --- | --- | --- |
|  |  | *Reg3y<sup>+/-</sup>* |  | *Reg3y<sup>-/-</sup>* |  | *Reg3y<sup>+/-</sup>* |  | *Reg3y<sup>-/-</sup>* |  |
|  |  | RC | HF | RC | HF | RC | HF | RC | HF |
| Actinomycetales | Micrococcaceae | 0 | 0 | 0 | 0 | 0 | 1 | 0 | 0 |
|  | Mycobacteriaceae | 1 | 0 | 0 | 0 | 0 | 0 | 0 | 0 |
| Bifidobacteriales | Bifidobacteriaceae | 1 | 0 | 0 | 0 | 0 | 1 | 0 | 1 |
| Coriobacteriales | Coriobacteriaceae | 0 | 0 | 0 | 1 | 0 | 0 | 0 | 0 |
| Bacteroidales | Bacteroidaceae | 0 | 0 | 0 | 0 | 0 | 0 | 1 | 0 |
|  | Porphyromonadaceae | 0 | 0 | 0 | 0 | 0 | 1 | 1 | 0 |
|  | Rikenellaceae | 1 | 0 | 1 | 1 | 0 | 0 | 1 | 0 |
|  | S24-7 | 4 | 6 | 7 | 0 | 7 | 3 | 8 | 6 |
| Deferribacterales | Deferribacteraceae | 1 | 0 | 0 | 0 | 0 | 0 | 0 | 0 |
| Streptophyta | Unassigned Streptophyta | 1 | 0 | 0 | 0 | 0 | 0 | 0 | 0 |
| Bacillales | Staphylococcaceae | 1 | 0 | 0 | 0 | 0 | 1 | 0 | 0 |
| Lactobacillales | Enterococcaceae | 0 | 0 | 0 | 0 | 0 | 1 | 0 | 0 |
|  | **Lactobacillaceae\*** | **2** | **0** | **3** | **4** | **1** | **1** | **3** | **3** |
|  | Streptococcaceae | 2 | 1 | 2 | 1 | 0 | 1 | 0 | 1 |
| Turicibacterales | Turicibacteraceae | 0 | 0 | 0 | 0 | 1 | 0 | 0 | 0 |
| Clostridiales | [Mogibacteriaceae] | 0 | 0 | 0 | 0 | 0 | 0 | 1 | 1 |
|  | **Clostridiaceae\*** | **1** | **2** | **1** | **1** | **0** | **1** | **1** | **1** |
|  | Lachnospiraceae | 0 | 0 | 0 | 2 | 0 | 3 | 6 | 5 |
|  | Peptococcaceae | 0 | 0 | 0 | 0 | 0 | 0 | 1 | 0 |
|  | **Peptostreptococcaceae\*** | **0** | **0** | **0** | **2** | **0** | **0** | **2** | **1** |
|  | Ruminococcaceae | 2 | 1 | 1 | 3 | 1 | 0 | 6 | 3 |
|  | Unassigned Clostridiales | 5 | 2 | 2 | 6 | 1 | 2 | 15 | 8 |
| Erysipelotrichales | Erysipelotrichaceae | 0 | 0 | 1 | 0 | 0 | 1 | 0 | 0 |
| Enterobacteriales | Enterobacteriaceae | 0 | 0 | 0 | 0 | 0 | 1 | 0 | 2 |
| Pseudomonadales | Pseudomonadaceae | 0 | 0 | 0 | 0 | 0 | 1 | 0 | 0 |
| Rhizobiales | Brucellaceae | 1 | 1 | 0 | 0 | 0 | 0 | 3 | 1 |
| Burkholderiales | Comamonadaceae | 0 | 0 | 0 | 0 | 0 | 0 | 0 | 1 |
|  | Oxalobacteraceae | 0 | 0 | 0 | 0 | 1 | 0 | 0 | 0 |
| CW040 | F16 | 0 | 1 | 0 | 0 | 0 | 0 | 1 | 0 |

\*Blue-shaded rows refer to families where relative abundances over a 12:12 LD period are presented in main text Figure 6, panel D.

